# Mechanical coupling between dorsal and ventral surfaces shapes the *Drosophila* haltere

**DOI:** 10.1101/2024.11.22.624869

**Authors:** Yuzhao Song, Paloma Martín, Tianhui Sun, Ernesto Sánchez-Herrero, José Carlos Pastor-Pareja

## Abstract

The extracellular matrix is an essential determinant of animal form, enabling organization of cells and tissues into organs with complex 3D shapes. In contrast with the dorso-ventrally flat *Drosophila* wing, its serial homolog the haltere adopts a globular shape thought to arise from a lack of matrix-mediated adhesion between its dorsal and ventral surfaces. Contradicting this model, however, matrix manipulations are known to deform halteres. To understand haltere morphogenesis, we characterized matrix behavior and monitored metamorphic development of the haltere. We found that, similar to the wing, correct haltere morphogenesis requires Collagen IV degradation, which we show is mediated by ecdysone-controlled expression of Matrix metalloprotease 2 in both wing and haltere. After Collagen IV is degraded, similar again to the wing, dorsal and ventral haltere surfaces establish Laminin-mediated contact through long cytoskeletal projections. Furthermore, time-lapse analysis of shape changes in wild type and mutant halteres indicates that these projections couple the two surfaces through a central tensor, ensuring load distribution across the whole organ to create a globular shape against tissue-wide deforming forces. Our findings reveal an unexpected role for matrix-mediated adhesion in haltere morphogenesis and describe a novel type of matrix-based tensor structure building 3D shape from 2D epithelia.

## INTRODUCTION

The epithelial tissue layer of apico-basally polarized cells is the fundamental architectural motif in the construction of animal body plans (Leys and Riesgo, 2012). Cells of an epithelium adhere to neighboring cells and attach basally to a specialized form of extracellular matrix (ECM) called basement membrane (BM). Despite their simple, two-dimensional nature, epithelia can be shaped into more complex 3D structures during development through different morphogenetic processes. Epithelial morphogenetic processes add or detract cells from the epithelium in particular regions, rearrange cells within the layer, change cell shape and height coordinately, or create folds by contraction/expansion of apical or basal cell surfaces (Bard, 1990; Fristrom, 1988; Schöck and Perrimon, 2002). Coherent transmission of forces and mechanical coupling of cells across the epithelium is ensured by the connection of each cell’s cytoskeleton, first, to neighboring cells through the cadherin-catenin complex and, second, to the BM through integrin-based focal adhesions (Lecuit et al., 2011). The cytoskeleton is additionally known to exert forces required in these morphogenetic changes, mainly through acto-myosin contractility and directional polymerization (Guillot and Lecuit, 2013; Heisenberg and Bellaiche, 2013). It is increasingly clear, however, that BMs play active morphogenetic roles as well, and that production, degradation and compositional changes in BMs are key generators of morphogenetic force in epithelia (Goodwin et al., 2016; Haigo and Bilder, 2011; Harmansa et al., 2023; Isabella and Horne-Badovinac, 2015; Khalilgharibi and Mao, 2021; Klussmann-Fricke et al., 2022; Ma et al., 2017; Matsubayashi et al., 2020; Pastor-Pareja and Xu, 2011; Ramos-Lewis and Page-McCaw, 2019; Wu et al., 2023), in coordination with the cytoskeleton or independently.

BMs are dense planar polymers of ECM glycoproteins and proteoglycans that basally underlie epithelia in all animals (Jayadev and Sherwood, 2017; Sekiguchi and Yamada, 2018; Yurchenco, 2011). Through the ECM receptor integrin and integrin-associated proteins, BMs are connected to the actin and microtubule epithelial cytoskeletons. BMs are made of four main conserved components: Laminin, Nidogen, Collagen IV and Perlecan (LNCP matrix). Despite the stereotypical planar morphology and conserved composition of most BMs, there is a growing number of documented examples of atypical BMs or BM-related matrices that are not planar or do not contain all four major components, and that are not assembly intermediates, but final structures with defined roles (Pastor-Pareja, 2020). In vertebrates, these include the BM plaques of the corneal stroma, the perichondrocyte matrix, the amorphous matrix of the mammalian trophoblast and double BMs in the kidney glomerulus and developing fins (Keeley and Sherwood, 2019; Pastor-Pareja, 2020). In the fruit fly *Drosophila melanogaster*, a particularly apt model for the genetic dissection of morphogenetic ECM roles, well studied examples of atypical BMs are the special matrix of the heart, the myotendinous junction in muscle attachment sites, and Collagen IV intercellular concentrations (CIVICs) in adipocytes (Dai et al., 2017; Pastor-Pareja, 2020). Other instances have been recently characterized in the lymph gland (Khadilkar et al., 2020), the fenestrated membrane of the eye (Ready and Chang, 2023; Walther et al., 2024) and wing dorso-ventral (DV) adhesion (Sun et al., 2021), the latter involving a Laminin-only matrix.

Insect wings are flat appendages formed by basal-to-basal adhesion between dorsal and ventral epithelial wing surfaces (Waddington, 1940). From genetic studies in *Drosophila* forewings, arising in the second thoracic (T2) segment, it is known that wing DV adhesion requires integrins and integrin-associated proteins (Prout et al., 1997; Walsh and Brown, 1998), as well as the BM component Laminin (Henchcliffe et al., 1993; Martin et al., 1999). However, because an intervening BM cannot be discerned from ultrastructural studies (Fristrom et al., 1993; Mogensen and Tucker, 1988), the mechanism of wing adhesion, taking place during metamorphosis, remained for a long time mysterious. It was later found that wing DV adhesion requires as its first step the degradation of the BMs underlying its dorsal and ventral surfaces (De las Heras et al., 2018; Diaz-de-la-Loza et al., 2018). Due to differential degradation of BM components and continued local Laminin production, these conventional BMs are replaced by a Laminin matrix to which both surfaces connect through long cytoskeletal projections containing both microtubules and F-actin (Mogensen and Tucker, 1988; Sun et al., 2021). These projections originate as a result of hydrostatic pressure that pushes the two Laminin-attached surfaces apart around the time of head eversion (12 h after puparium formation [APF]), and their correct formation requires microtubule-binding proteins Short stop (Shot) and SAXO downstream of blistered (Sdb) (Sun et al., 2021). Later, DV projections disassemble and contract, dramatically shortening the distance between the two surfaces and flattening the wing (Singh et al., 2024; Sun et al., 2021; Tran et al., 2024). After DV adhesion, other morphogenetic events contribute to shaping the wing during metamorphosis: hinge contraction, which exerts a proximal pull on the whole tissue (Aigouy et al., 2010; Etournay et al., 2015), anchorage of the wing to the margin cuticle (Ray et al., 2015), localized apoptosis in the anterior-proximal region (Matamoro-Vidal et al., 2024), and wing expansion/folding (Tsuboi et al., 2023).

In contrast with the flat forewing in the second thoracic (T2) segment, in *Drosophila* and all dipteran insects (flies and mosquitoes), the T3 hindwing adopts the form of a globular, club-like organ called “haltere”. Dipteran halteres serve no aerodynamic purpose, but are essential for flight control and steering, and responsible for the unparalleled maneuvering displayed by this order of insects (Dickerson et al., 2019). Despite their strikingly divergent adult morphology, the T3 haltere and the T2 wing develop from similar imaginal discs, patterned through almost identical mechanisms, albeit the haltere disc is smaller. Differential haltere development is controlled by the Hox gene *Ultrabithorax (Ubx)*, loss of which homeotically transforms halteres into wings (Lewis, 1978). While size difference between the haltere and the wing is understood and involves larval Dpp gradient modulation by Ubx (Crickmore and Mann, 2006; de Navas et al., 2006; Weatherbee et al., 1998), causes for the difference in overall morphology, arising later during metamorphosis, are less clear. According to one theory, the critical difference between the flat wing and the globular haltere is the lack of DV adhesion between dorsal and ventral haltere surfaces. Absence of DV adhesion in the haltere would be determined by a lack of BM degradation, which in the wing takes place between 4 and 7 h APF (De las Heras et al., 2018; Diaz-de-la-Loza et al., 2020; Diaz-de-la-Loza et al., 2018). In this differential degradation scenario, the haltere, because of its lack of BM degradation, does not undergo initial DV contact and hence its surfaces stay separated throughout the remainder of the pupal period. Against this “hollow haltere” theory, however, overexpression of matrix metalloprotease inhibitor TIMP (Tissue Inhibitor of Metallo-Proteases) impairs not just development of the wing, but also of the haltere (De las Heras et al., 2018), raising the possibility that BM degradation plays a role in haltere morphogenesis as well. Furthermore, histological sections of the pupal halteres do not show a hollow appendage, but a structure filled with cellular material of unknown organization and origin (Roch and Akam, 2000). Therefore, key aspects of the metamorphic development of the haltere and its morphological difference with the wing remain to be elucidated.

In this study, we aimed at understanding the role of the BM in the morphogenesis of the haltere. Through genetic manipulation and in vivo imaging, we show that early BM degradation and DV adhesion do take place in the haltere, as in the wing, and that BM degradation requires Mmp2 in both appendages. After BM degradation, a Laminin matrix remains to which long projections from dorsal and ventral haltere surfaces connect. Unlike wing DV projections, haltere DV projections do not subsequently contract, leaving the haltere unflattened. Importantly, we found that DV contact mediated by DV projections counters opposing deforming forces exerted by the contraction of the proximal haltere and the attachment of its distal region to the cuticle. Therefore, DV adhesion occurs in the haltere and is essential for it to acquire a globular shape.

## RESULTS

### Haltere BMs are degraded during early metamorphosis

The adult haltere, derived from dorsal and ventral epithelial regions of the haltere larval imaginal disc, acquires its globular, club-like shape during metamorphosis (Figure 1A). This shape contrasts with the flat adult wing, which develops from a larger but otherwise very similar imaginal disc. To understand the metamorphic development of the haltere, we first analyzed the presence of BM components between the dorsal and ventral surfaces of the haltere during the prepupal period up to head eversion (12 h APF), marking the beginning of true metamorphosis (prepupa-to-pupa transition). To do that, we imaged functional fluorescently tagged transgenic versions of the four main BM proteins expressed under control of their promoters: Collagen IV (α2 chain Vkg), Perlecan (Trol), Nidogen (Ndg) and Laminin (β chain LanB1). Previous studies reported that Collagen IV is not degraded in the haltere, contrasting with its decrease in the wing, where it is degraded between 4 and 7 h APF (De las Heras et al., 2018; Diaz-de-la-Loza et al., 2020; Diaz-de-la-Loza et al., 2018). When we examined Collagen IV presence in prepupal halteres at 8 and 12 h APF, however, we observed a sharp decrease (Figure 1B). Furthermore, the amounts of Perlecan (Figure 1C) and Nidogen (Figure 1D) were similarly decreased. Laminin, albeit reduced, suffered the least decrease of all four main BM components (Figure 1E), reminiscent of our previous findings in the wing (Sun et al., 2021). Altogether, these observations suggest that the BMs underlying the dorsal and ventral surfaces of the prepupal haltere undergo a degradation process similar to the prepupal wing, albeit with a slight delay.

**Figure 1.**
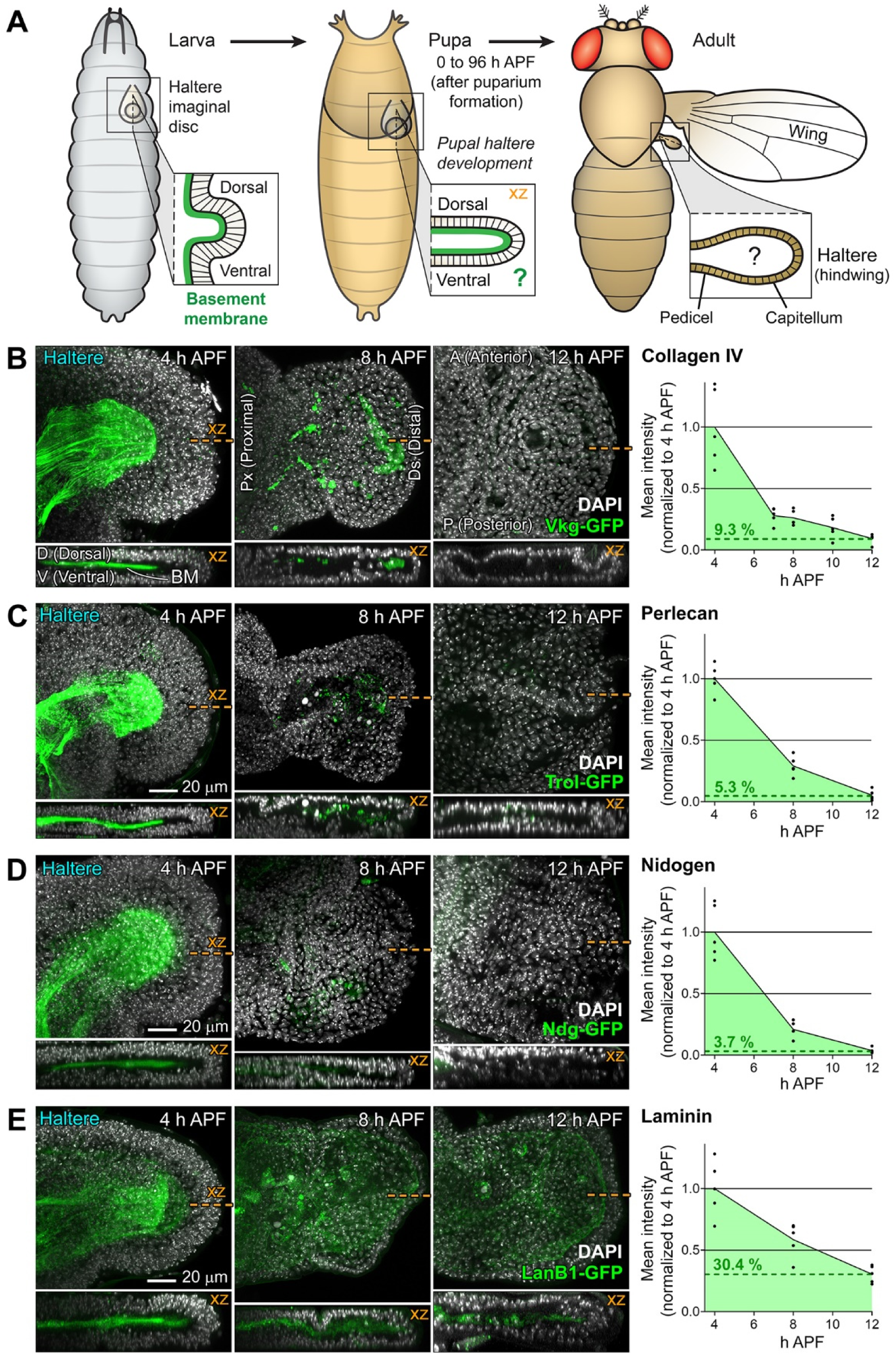
Haltere BMs are degraded during early metamorphosis. (A) Metamorphic development of the adult haltere from its corresponding imaginal disc in the larva. (B) Prepupal halteres dissected at 4, 8 and 12 h APF from flies endogenously expressing a functional GFP-tagged version of BM component Collagen IV (Vkg^G454^-GFP α2 chain; green). Images are maximum intensity projections (MIP) of confocal xy sections. Anterior [A] up, posterior [P] down; proximal [Px] left, distal [Ds] right. Nuclei stained with DAPI (white). Lower panels show xz sections (dorsal [D] up, ventral [V] down) of the same halteres at positions indicated by discontinuous orange lines. The graph represents individual haltere measurements of mean GFP intensity (intensity divided per tissue area) and average from 5 haltere images per time point. Values are normalized to the average intensity at 4 h APF. Dotted lines indicate 12 h APF GFP intensity as a percentage of 4 h APF intensity. (C) Halteres dissected at 4, 8 and 12 h APF from flies endogenously expressing a functional GFP-tagged version of BM component Perlecan (Trol^ZCL1700^-GFP, green). Lower panels show xz sections. The graph represents measurements of GFP intensity as in (B). (D) Halteres dissected at 4, 8 and 12 h APF from flies expressing under control of its own promoter a functional GFP-tagged version of BM component Nidogen (Ndg^fTRG.638^-sGFP, green). Lower panels show xz sections. The graph represents measurements of GFP intensity as in (B). (E) Halteres dissected at 4, 8 and 12 h APF from flies expressing under control of its own promoter a functional GFP-tagged version of BM component Laminin (LanB1^fTRG.631^-sGFP β chain, green). Lower panels show xz sections. The graph represents measurements of GFP intensity as in (B).

### Mmp2 expression downstream of ecdysone degrades BM Collagen IV in both haltere and wing

Having observed decreased presence of BM components in the prepupal haltere, we sought to understand its regulation and effects on haltere morphogenesis. To do that, we manipulated the activity of matrix metalloproteinases (MMPs), a family of conserved proteases involved in ECM clearance in multiple developmental contexts (Page-McCaw et al., 2007). The *Drosophila* genome encodes two MMPs, namely Mmp1 and Mmp2 (Page-McCaw et al., 2003), and the conserved MMP inhibitor TIMP (Godenschwege et al., 2000). TIMP overexpression and Mmp2 knock down under control of *rn-GAL4* caused Collagen IV accumulation between dorsal and ventral haltere surfaces, observed at both 8 h APF (Figure 2A and B) and 12 h APF (Figure 2C and D). We found a similar accumulation of Collagen IV upon knock down in the haltere of the Ecdysone Receptor (EcR) (Figure 2A-D), responsible for transducing signaling by the steroid hormone ecdysone and coordinating multiple morphogenetic events during metamorphosis (Yamanaka et al., 2013). Furthermore, overexpression of TIMP and knock down of EcR or Mmp2 (but not Mmp1; Suppl. Table S1) impaired haltere morphogenesis and produced rounder appendages, whereas Mmp2 overexpression resulted in elongated halteres (Figure 2E). Knock down of Mmp2 or EcR also prevented Collagen IV degradation in the wing (Figure 2F-H), leading to blisters (Figure 2I), as previously shown to occur upon TIMP overexpression (De las Heras et al., 2018; Diaz-de-la-Loza et al., 2018; Sun et al., 2021). In both haltere and wing, knock down of EcR under *rn-GAL4* control reduced Mmp2 expression, as measured by qRT-PCR (Figure 2J). Furthermore, *Mmp2* expression was upregulated in the whole animal during the larva-to-prepupa transition (Figure 2K), consistent with *Mmp2* being a general ecdysone-regulated gene. Altogether, these data indicate that *Mmp2* expression is required downstream of EcR signaling for Collagen IV degradation in both haltere and wing (Figure 2L). Importantly, in the haltere context, these results demonstrate that degradation of BM components takes place in the haltere and is required for its correct morphogenesis.

**Figure 2.**
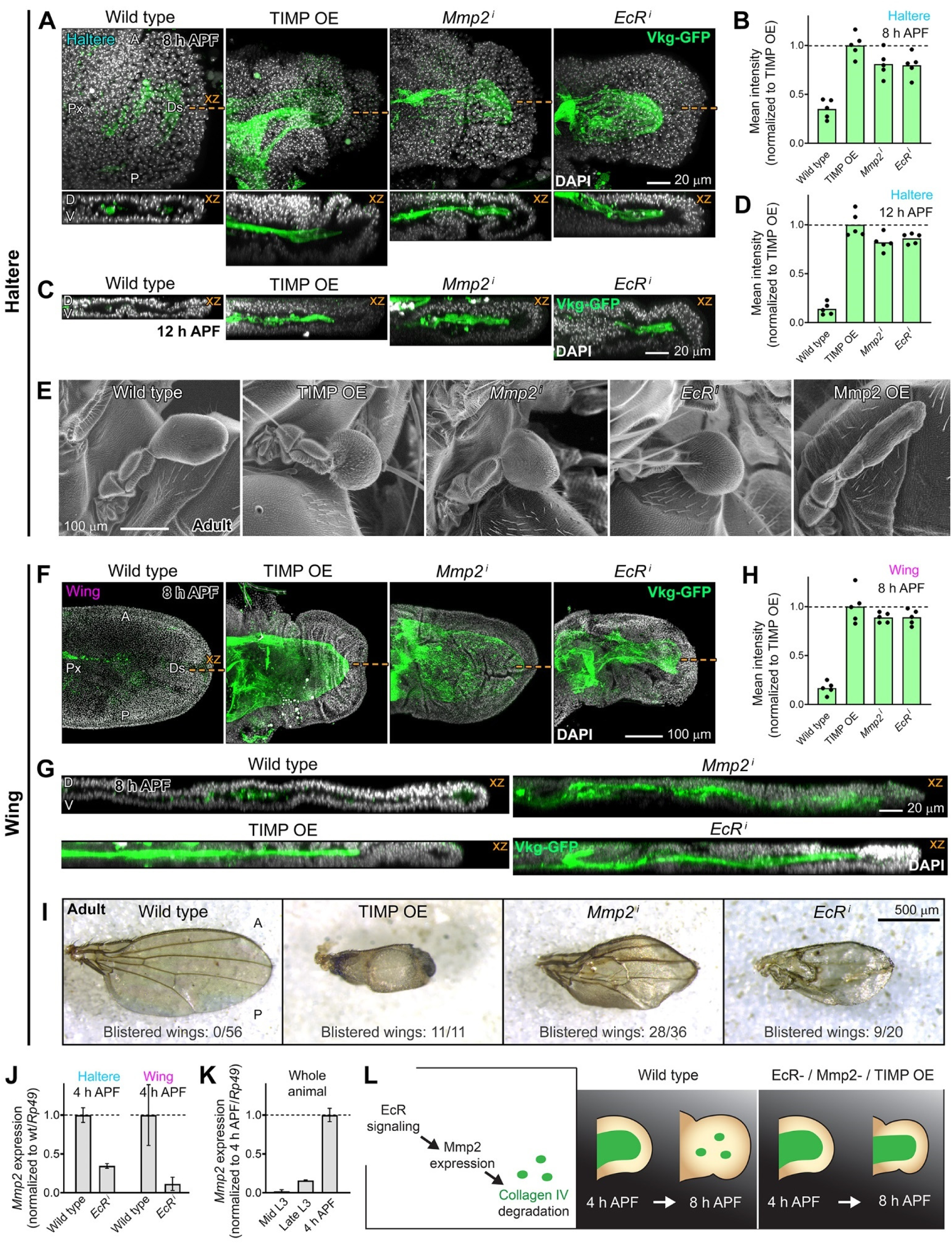
Mmp2 expression downstream of ecdysone degrades BM Collagen IV in both haltere and wing. (A) Images (MIP of confocal xy sections) of a wild type prepupal haltere and halteres where MMP inhibitor TIMP was overexpressed (*rn>*TIMP OE), or Mmp2 (*rn>Mmp2^i^*) and Ecdysone Receptor (*rn>EcR^i^*) knocked down under control of *rn-GAL4*. Halteres were dissected at 8 h APF from flies endogenously expressing a functional GFP-tagged version of Collagen IV (Vkg^G454^-GFP, green). Images are maximum intensity projections of xy sections. Anterior [A] up, posterior [P] down; proximal [Px] left, distal [Ds] right. Nuclei stained with DAPI (white). Lower panels show xz sections (dorsal [D] up, ventral [V] down) of the same halteres at positions indicated by discontinuous orange lines. (B) Vkg-GFP mean intensity (total intensity divided per tissue area) measured in wild type, TIMP OE, *Mmp2^i^* and *EcR^i^* halteres dissected at 8 h APF. Values are normalized to the average TIMP OE intensity. Bar height represents average value from 5 measurements per genotype. (C) Confocal xz sections of wild type, TIMP OE, *Mmp2^i^* and *EcR^i^* halteres dissected at 12 h APF. Vkg-GFP in green. Nuclei in white (DAPI). (D) Vkg-GFP mean intensity in wild type, TIMP OE, *Mmp2^i^* and *EcR^i^* halteres at 12 h APF as in (B). (E) Scanning electron micrographs of wild type, TIMP OE, *Mmp2^i^*, *EcR^i^* and Mmp2 OE adult halteres. (F) Images (MIP of confocal xy sections) of wild type, *rn>*TIMP OE, *rn>Mmp2^i^* and *rn>EcR^i^* prepupal wings dissected at 8 h APF. Anterior [A] up, posterior [P] down; proximal [Px] left, distal [Ds] right. Vkg-GFP in green. Nuclei stained with DAPI (white). (G) Confocal xz sections (dorsal [D] up, ventral [V] down) of the wild type, TIMP OE, *Mmp2^i^* and *EcR^i^* wings in (F), dissected at 8 h APF. Vkg-GFP in green. Nuclei in white (DAPI). (H) Vkg-GFP mean intensity in wild type, TIMP OE, *Mmp2^i^* and *EcR^i^* wings at 8 h APF as in (B). (I) Wild type, TIMP OE, *Mmp2^i^* and *EcR^i^* adult wings. Penetrance of blistering in each genotype indicated as the fraction of blistered wings per wings examined. (J) *Mmp2* expression by qRT-PCR in wild type and *EcR^i^* wings and halteres at 4 h APF. Expression levels are normalized to wild type levels and *rp49*. Error bars represent SD (n=3). (K) *Mmp2* expression by qRT-PCR in whole wild type mid L3 larvae, late L3 larvae and 4 h APF prepupae. Expression levels are normalized to 4 h APF levels and *rp49*. Error bars represent SD (n=3). (L) Mechanism of BM degradation in the prepupal haltere (and wing), and the effect of EcR knock down, Mmp2 knock down and TIMP overexpression, preventing Collagen IV clearance.

### Microtubule-actin projections mediate DV adhesion in the haltere

To elucidate the role of the ECM in the morphogenesis of the haltere during metamorphosis, we imaged halteres at different times after head eversion. In order to preserve normal tissue morphology, we conducted confocal imaging in live animals (see Methods), as initial attempts to image fixed halteres in flat preparations did not allow us to discern their 3D architecture and key morphological features. At 14 h APF (2 h after head eversion [AHE]), we observed that the dorsal and ventral surfaces of the haltere were connected by long basal cell projections containing both microtubules and F-actin, labeled with Tubulin-GFP and Lifeact-RFP, respectively (Figure 3A). The main region of DV contact mediated by these projections formed a proximo-distal stripe along the anterior haltere, whereas a minor contact area was present in a posterior, distal position. At 24 h APF, the haltere presented a more compact shape, with the DV contact area reduced to a central spot consisting of shorter projections (Figure 3B). At 40 h APF, the haltere had compacted further and displayed a morphology akin to the adult haltere, with a distinct globular capitellum and constricted pedicel (Figure 3C). At this time, the center of the capitellum was occupied by a tongue-shaped structure to which projections from both haltere surfaces connected. This DV contact structure concentrated Laminin (Figure 3D), the ECM receptor integrin (Figure 3E) and integrin-associated Talin (Figure 3F), suggesting an accumulation of focal adhesions. Furthermore, we found that the central tongue-shaped structure was surrounded by a tubular channel (Figure 3G), similar to the way DV-ahered intervein regions in the wing are delimited by non-adhered veins and margins (Fristrom et al., 1994). In addition to this channel, within each haltere surface, there were abundant interstitial spaces left between basal projections, reminiscent of the Parthenon-like structure recently described in the pupal leg (Hiraiwa et al., 2024). In summary, our characterization of haltere morphogenesis shows that adhesion between its dorsal and ventral surfaces are defining features of its metamorphic development (Figure 3H). Together with evidence for an Mmp2 requirement and persistence of Laminin, our results reveal that the metamorphic development of the haltere surprisingly resembles that of the wing in key aspects, with BM degradation and DV adhesion taking place in the haltere as well.

**Figure 3.**
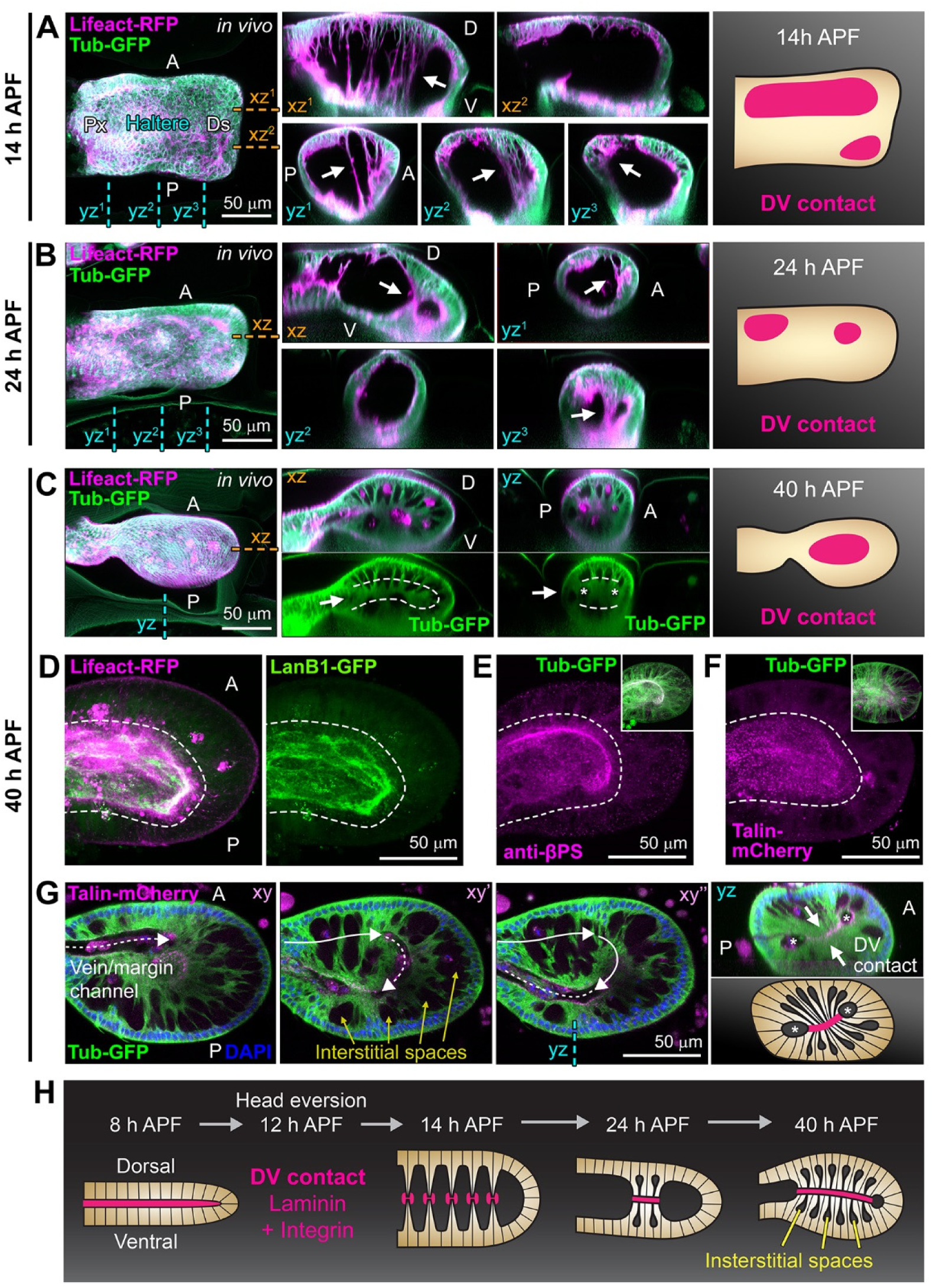
Microtubule-actin projections mediate DV adhesion in the haltere. (A) In vivo image (MIP of confocal dorsal xy sections) of a pupal haltere (14 h APF = 2 h AHE) where F-actin and microtubules are labeled by expression of Lifeact-RFP (magenta) and Tub-GFP (α-Tubulin 84B, green) under *rn-GAL4* control. Anterior [A] up, posterior [P] down; proximal [Px] left, distal [Ds] right. Position of xz and yz sections indicated by discontinuous orange and cyan lines, respectively. Arrows point to DV projections. The drawing on the right depicts DV contact areas. (B) In vivo image of a pupal haltere (24 h APF = 12 h AHE) and drawing of DV contact areas as in (A). Arrows point to DV contacts. (C) In vivo image of a pupal haltere (40 h APF) and drawing of DV contact areas as in (A). Arrows point to the DV contact area, outlined by dashed lines. Asterisks mark the surrounding tubular channel. (D) Confocal section of an haltere dissected at 40 h APF from a pupa expressing a functional GFP-tagged version of Laminin (LanB1^fTRG.631^-sGFP, green) under control of its own promoter and *rn-GAL4*-driven Lifeact-RFP (magenta). The image on the right shows LanB1-GFP signal separately. DV contact area outlined by a dashed line. (E) Confocal section of an haltere dissected at 40 h APF from a pupa stained with an antibody against the integrin β subunit Mys (anti-βPS; magenta) and *rn-GAL4*-driven Tub-GFP (green). The inset shows both signals together and the larger image anti-βPS separately. DV contact area outlined by a dashed line. (F) Confocal section of an haltere dissected at 40 h APF from a pupa expressing a functional mCherry-tagged version of Talin (Rhea^MI00296-mCh.0^-mCherry, magenta) endogenously and *rn-GAL4*-driven Tub-GFP (green). The inset shows both signals together and the larger image Talin-mCherry separately. DV contact area outlined by a dashed line. (G) Confocal sections (xy to xy’’: dorsal to ventral) of an haltere dissected at 40 h APF from a pupa expressing Talin-mCherry (magenta) endogenously and *rn-GAL4*-driven Tub-GFP (green). Yellow arrows point to interstitial spaces. White arrows in xy sections follow the tubular channel surrounding DV contact area. On the yz section, arrows point to DV contact (notice Talin-mCherry line) and asterisks mark the tubular channel. (H) Changes in haltere architecture up to 40 h APF. At 8 h APF, after degradation of dorsal and ventral BMs, both surfaces are connected by a Laminin matrix. Next, DV contact is maintained through DV projections, probably formed, like in the wing, as a result of increased hydrostatic pressure that pushes surfaces apart around head eversion (12 h APF). Later, the haltere undergoes compaction while projections coalesce into a central DV contact area.

Having shown that DV adhesion is not an exclusive feature of wing development, we aimed to ascertain when the morphological difference between the flat wing and the globular haltere arises. For that, we recorded the development of halteres and wings expressing Tub-GFP and Lifeact-RFP after head eversion (Suppl. Movie S1). Between 14 and 26 h APF (2 to 14 h AHE), we observed a gradual compaction of the haltere and the coalescence of the anterior stripe of DV projections into a central contact spot (Figure 4A). As for the wing, we confirmed our previous observations (Sun et al., 2021) that flattening coincides with the sudden disassembly of DV projections between 20 and 24 h APF (Figure 4B). Quantitation of cell behaviors in these movies revealed remarkable differences between haltere and wing cells. First, the apical area of haltere cells kept gradually shrinking during the imaging period; wing apical areas, in contrast, underwent an initial increase, consistent with a recent report (Singh et al., 2024), but then rapidly decreased to approximately half of their previous maximum (Figure 4C). Flattening of the organ, measured as distance from the apical surface to the DV plane, experienced in the haltere an initial drop to become rather stable around 35 μm at 18 h APF; in wing cells, in contrast, sudden disassembly of DV projections brought epithelial height, initially similar to the haltere, to approximately 15 μm (Figure 4D). A similar contrast between gradual haltere changes and faster ones in the wing was observed in the detachment from the apical cuticle, to which both initially adhered: distancing of haltere cells from the cuticle was progressive, whereas wing cells detached more abruptly (Figure 4E). These results indicate that differences between haltere and wing in cell shape and epithelial height are not dictated by DV adhesion in the haltere, since both haltere and wing display DV contact, but are cell-intrinsic. Supporting this, in *postbithorax* (*pbx*) flies (*pbx^1^*/Df *Ubx^109^*), where a regulatory *Ultrabithorax* (*Ubx*) mutation homeotically transforms the posterior compartment of the haltere into wing (Bender et al., 1983) (Figure 4F), DV contact exists at 40 h APF in both the haltere and wing parts of the mutant appendage despite the difference in epithelial height (Figure 4G). In all, our analysis demonstrates that DV adhesion is not a distinguishing feature between haltere and wing. Instead, the differential behavior determining that the haltere remains globular while the wing becomes flat is the fast shortening of DV projections at 20-24 h APF, taking place in the wing but not the haltere.

**Figure 4.**
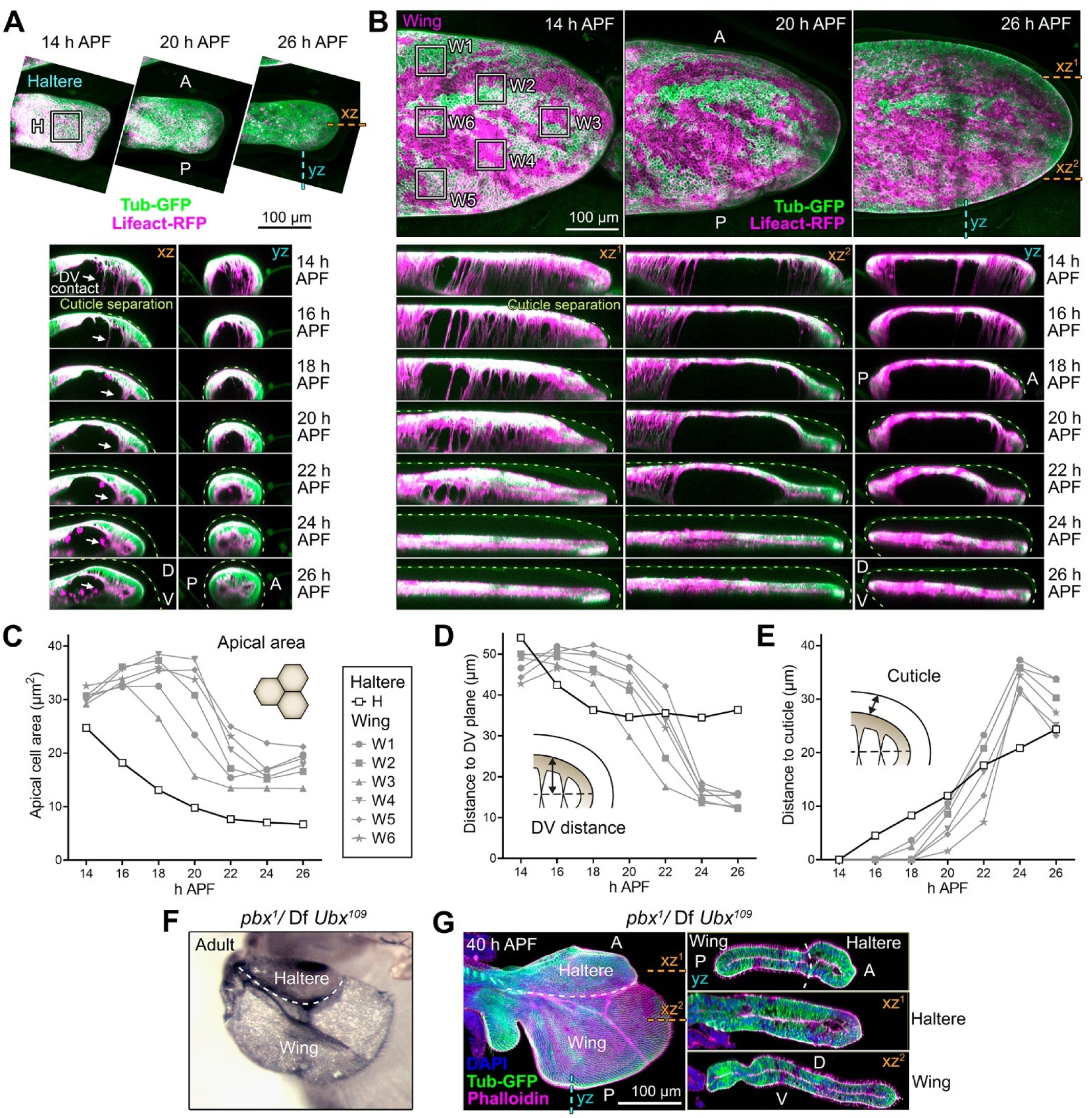
Comparative analysis of cell and tissue changes in haltere and wing after head eversion. (A) Time-lapse imaging (MIP of xy confocal dorsal sections) of haltere development from 14 to 26 h APF (2 to 14 h AHE). Expression of Lifeact-RFP (F-actin, magenta) and Tubulin-GFP (microtubules, green) driven by *rn-GAL4*. Anterior [A] up, posterior [P] down. Position of xz and yz sections below (dorsal [D] up, ventral [V] down) indicated by discontinuous orange and cyan lines, respectively. Dashed green lines in xz and yz sections mark the chitinaceous cuticle, visible due to autofluorescence. Arrows point to DV projections. See Suppl. Movie S1. (B) Time-lapse imaging (MIP of xy confocal dorsal sections) of wing development from 14 to 26 h APF. F-actin (magenta) and microtubules (green) labeled by expression of Lifeact-RFP and Tub-GFP, respectively. Anterior [A] up, posterior [P] down. Position of xz and yz sections below (dorsal [D] up, ventral [V] down) indicated by discontinuous orange and cyan lines, respectively. Dashed green lines in xz and yz sections mark the cuticle. See Suppl. Movie S1. (C) Changes in apical cell area in the haltere central region (H square in (A)) and different wing regions (W1-6 squares in (B)). Points represent average values of dorsal cells inside the corresponding square at indicated times. (D) Changes in DV distance in the haltere central region and different wing regions. Points represent distance from the apical dorsal surface to the basal DV contact in the center of the corresponding square. (E) Changes in distance to cuticle in the haltere central region and different wing regions. Points represent distance from the apical dorsal surface to the cuticle in the center of the corresponding square. (F) Adult haltere in a *postbithorax* mutant (*Ubx* regulatory mutant allele *pbx^1^*over a deficiency uncovering *Ubx*). The posterior compartment of the haltere is homeotically transformed into wing. The dashed line separates anterior haltere and posterior (transformed) wing tissue. (G) Confocal image (MIP of xy confocal sections) of a *postbithorax* haltere dissected at 40 h APF. <u>F-actin</u> <u>(magenta) and microtubules (green) labeled by phalloidin and</u> *rn-GAL4*-driven <u>Tub-GFP, respectively</u>. The dashed line separates haltere and wing tissue. Position of yz and xz sections on the right (dorsal [D] up, ventral [V] down) indicated by discontinuous cyan and orange dashed lines. Note DV distance is higher in the haltere part.

### Correct haltere morphogenesis requires Laminin-dependent DV adhesion and microtubule-binding proteins Shot and Sdb

Once established that the haltere displays DV adhesion through a central structure containing Laminin and integrin, we decided to study the role of DV contact and cytoskeletal projections in haltere morphogenesis. We previously showed that Laminin and integrin are required for the formation of DV microtubule-actin projections in the wing, and that microtubule-binding proteins encoded by *shot* and *Sdb* are essential to make those projections resistant to the hydrostatic pressure that pushes the wing surfaces apart during head eversion (Sun et al., 2021). We therefore knocked down the expression in the haltere of laminin (common β subunit of the Laminin heterotrimer LanB1), integrin (β subunit of the integrin receptor Mys) and the integrin-associated cytoskeletal linker talin (Rhea). In contrast with the wild type (Figure 5A), DV projections were absent at 14 h APF in *LanB1^i^*, *mys^i^* and *rhea^i^* halteres (Figure 5B). DV projections, however, were present when we knocked down Shot and Sdb (Figure 5C), suggesting that, similar to the wing, they are not required to form projections in the first place (Sun et al., 2021). Later, at 40 h APF, *LanB1^i^*, *mys^i^* and *rhea^i^*halteres still lacked DV projections and the central DV contact structure had not formed (Figure 5D and E). Strikingly, the morphology of these halteres differed from the wild type: instead of globular, they appeared elongated in the proximo-distal axis, lanceolated in shape and flatter in section. We observed similar deformations in *shot^i^* and *Sdb^i^* halteres (Figure 5F), suggesting that DV projections were present but defective. Furthermore, mutant haltere shapes at 40 h APF resembled those of the corresponding adults (Figure 5G-I). We previously showed that Sdb is expressed in the wing under control of transcription factor SRF (Serum Response Factor), encoded by *blistered* (*bs*) (Sun et al., 2021). In the haltere, we found that a *bs-GAL4* reporter was expressed at 40 h APF in the center of the capitellum region in both dorsal and ventral surfaces (Figure 5J). Moreover, anti-Sdb staining showed that Sdb was expressed in this same pattern and depended on *bs*, as *bs* knock down abrogated Sdb expression (Figure 5K and L), producing elongated halteres (Figure 5M). These data show that, similar to the wing, correct haltere morphogenesis requires focal adhesion-dependent formation of DV projections containing Shot and Sdb. Intriguingly, the loss of these factors in the wing causes blisters, a defect with no obvious parallel with the elongation of halteres. A final condition we found affected haltere shape was loss of *dumpy* (*dpy*), encoding a giant zona pellucida protein that maintains apical attachment of wing and legs to cuticle (Ray et al., 2015; Wilkin et al., 2000). *dpy^i^*halteres were shorter in the proximo-distal axis and rounder in shape than the wild type (Figure 5N), suggesting a process in which cuticle attachment and DV adhesion exert opposing organ-shaping forces.

**Figure 5.**
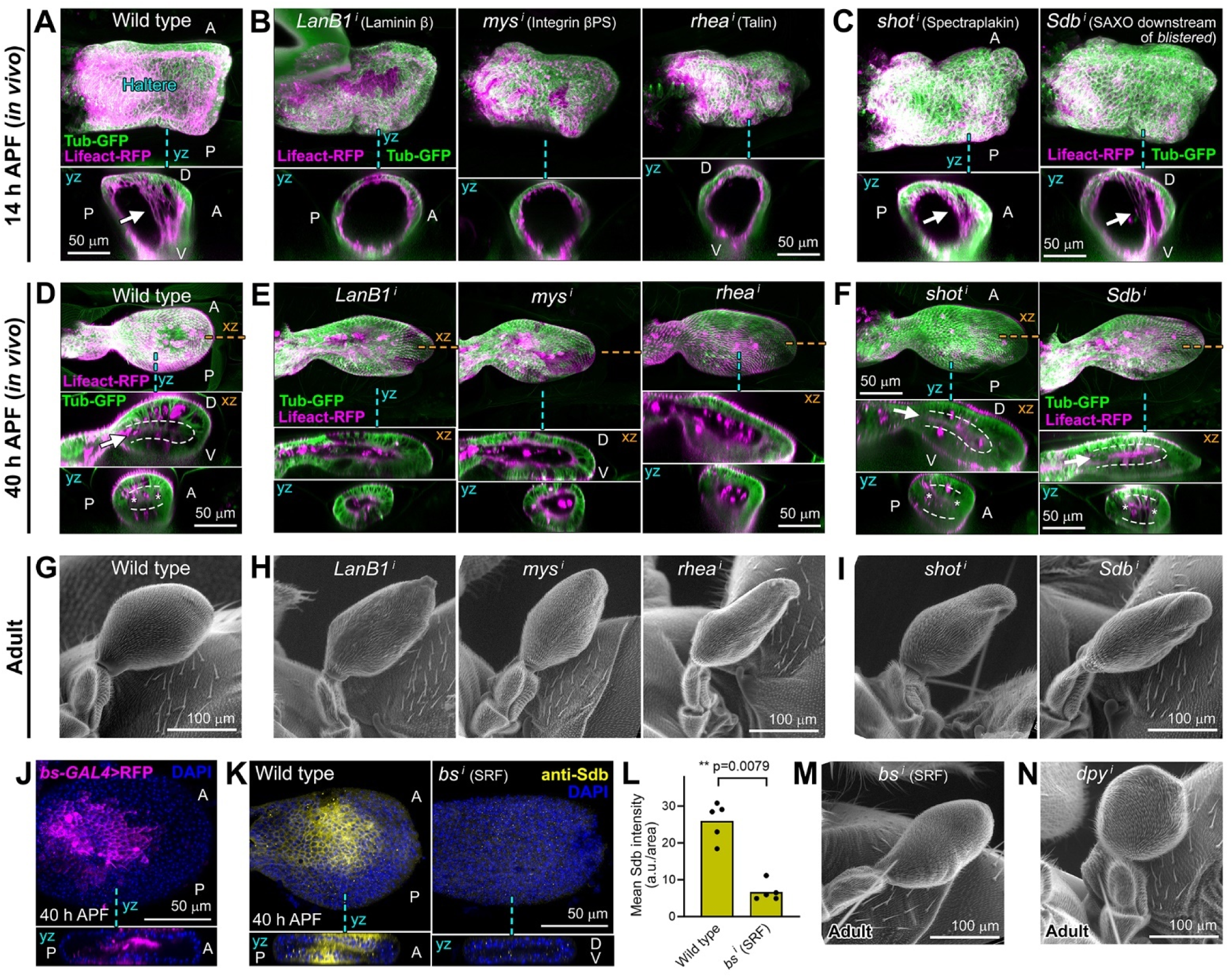
Correct haltere morphogenesis requires Laminin-dependent DV adhesion and microtubule-binding proteins Shot and Sdb. (A) In vivo image (MIP of confocal dorsal xy sections) of a wild type pupal haltere at 14 h APF. F-actin and microtubules were labeled by expression of Lifeact-RFP (magenta) and Tubulin-GFP (green) driven by *rn-GAL4*, respectively. Anterior [A] up, posterior [P] down. Position of yz section below indicated by a discontinuous cyan line. Arrow points to DV projections. (B) In vivo image of *rn>LanB1^i^* (Laminin β knock down), *rn>mys^i^* (Integrin βPS knock down) and *rn>rhea^i^* (talin knock down) halteres at 14 h APF. Lifeact-RFP in magenta; Tub-GFP in green. Note hollow lumen and absence of DV projections. (C) In vivo image of *rn>shot^i^* and *rn>Sdb^i^* halteres at 14 h APF. Lifeact-RFP in magenta; Tub-GFP in green. Arrow points to DV projections. (D) In vivo images (MIP of dorsal confocal xy sections) of a wild type pupal haltere at 40 h APF. F-actin (magenta) and microtubules (green) labeled by expression of Lifeact-RFP and Tub-GFP, respectively. Position of xz and yz sections below indicated by discontinuous orange and cyan lines, respectively. Arrow points to the DV contact area, outlined by white dashed lines, and asterisks mark the surrounding channel. (E) In vivo images of *LanB1^i^*, *mys^i^* and *rhea^i^*halteres at 40 h APF. Lifeact-RFP in magenta; Tub-GFP in green. Note hollow lumen, absence of DV projections and elongated shape. (F) In vivo images of *shot^i^* and *Sdb^i^* halteres at 40 h APF. Lifeact-RFP in magenta; Tub-GFP in green. Arrow points to DV contact area, outlined by white dashed lines, and asterisks mark the surrounding. Note elongated shape. (G) Scanning electron micrographs of a wild type adult haltere. (H) Scanning electron micrographs of *LanB1^i^*, *mys^i^* and *rhea^i^* adult halteres. Note elongated shape. (I) Scanning electron micrographs of *shot^i^* and *Sdb^i^* adult halteres. Note elongated shape. (J) Confocal image of a wild type haltere dissected at 40 h APF, expressing Lifeact-RFP (magenta) under control of *bs-GAL4* dissected at 40 h APF. Position of yz section below indicated by a discontinuous cyan line. Nuclei stained with DAPI (blue). (K) Confocal images of wild type and *rn>bs^i^* halteres dissected at 40 h APF, stained with anti-Sdb (yellow). Position of yz sections below indicated by discontinuous cyan lines. Nuclei stained with DAPI (blue). (L) Sdb antibody staining intensity measured in 5 images like those in (K) per genotype. Difference between wild type and *bs^i^* is significant in a Mann-Whitney test (p<0.01, **). (M) Scanning electron micrograph of a *bs^i^* adult haltere. Note elongated shape. (N) Scanning electron micrograph of a *rn>dpy^i^* (knock down of apical ECM protein Dumpy) adult haltere. Note rounded shape.

### DV adhesion shapes the haltere by opposing tissue-wide deforming forces

To understand the role of DV adhesion in shaping the haltere, we recorded from 30 to 40 h APF the development of wild type, *rhea^i^* (no DV adhesion), *shot^i^* (defective DV projections) and *dpy^i^*(no attachment to cuticle) halteres. We used endogenous GFP-tagged septate junction protein Neurexin IV (NrxIV) to image cell outlines in three movies of the dorsal surface per genotype (Figure 6A; Suppl. Figure S1A-D; Suppl. Movie S2). Overall, our movies showed gradual elongation of *rhea^i^*and *shot^i^* halteres, whereas wild type halteres maintained a lenticular shape and *dpy^i^* halteres became rounder as they increased their distance from the distal cuticle and retracted proximally (Suppl. Movie S2; Suppl. Figure S1E). We analyzed movies using Epitools Icy software (Heller et al., 2016) to quantify changes in cell shape and apical area (see Methods), and help determine the tissue-shaping forces at play. In wild type and all three mutants, our movies showed strong constriction of the pedicel, where both contraction of individual cells and basal extrusions could be observed (Figure 6B; Suppl. Movie S2). In the capitellum, total area decreased in wild type halteres (−8.9% ± 6.1 SD), whereas *dpy^i^* halteres experienced an even deeper area decrease (−31.3% ± 8.3 SD), and *rhea^i^*and *shot^i^* halteres increased their area instead of decreasing it (14.3 % ± 2.5 SD and 22.8% ± 4.4 SD, respectively) (Figure 6C). When we analyzed individual cell areas, we found that average apical cell area changes closely approximated total tissue values: −8.2% ± 5.8 SD average change in wild type, −24.5% ± 5.3 SD in *dpy^i^*, 16.8 % ± 5.3 SD for *rhea^i^* and and 31.1% ± 8.3 SD for *shot^i^* (Figure 6D). This suggests that cell shape changes accounted well for tissue area changes, while contributions from other cellular processes may be minor. Apical cell area changes were not homogeneously distributed along the proximo-distal axis, with changes in the proximal region contributing more to the compaction of wild type and *dpy^i^* halteres, and the distal part experiencing most of the expansion in the *rhea^i^* and *shot^i^* halteres (Figure 6E). Furthermore, distal cell area increases in *rhea^i^* and *shot^i^* halteres were directional, with the proximo-distal component of their deformation surpassing the antero-posterior one (Figure 6F; Suppl. Figure S1F). In all, in vivo imaging analysis indicates that the organ is shaped by the following forces (Figure 6G): a contraction of the pedicel that pulls the capitellum proximally; a reaction to the pedicel pull dictated by the attachment to the distal cuticle; balancing these two opposing forces, projections connected through DV adhesion act as a tensor that absorbs and distributes proximal and distal pulls, preventing their exclusive transmission through the surface and elongation observed in *rhea^i^* and *shot^i^* halteres (Figure 6E and F). In addition to these global forces, a fourth factor shaping the haltere is autonomous cell contraction in the capitellum, stronger in proximal regions, as evidenced by the proximal area decrease in wild type and *dpy^i^* halteres (Figure 6E and F). In summary, DV contact in the haltere acts a tensor that preserves a compact, globular shape and prevents deformation against opposing pulls from the proximal pedicel and distally attached cuticle.

**Figure 6.**
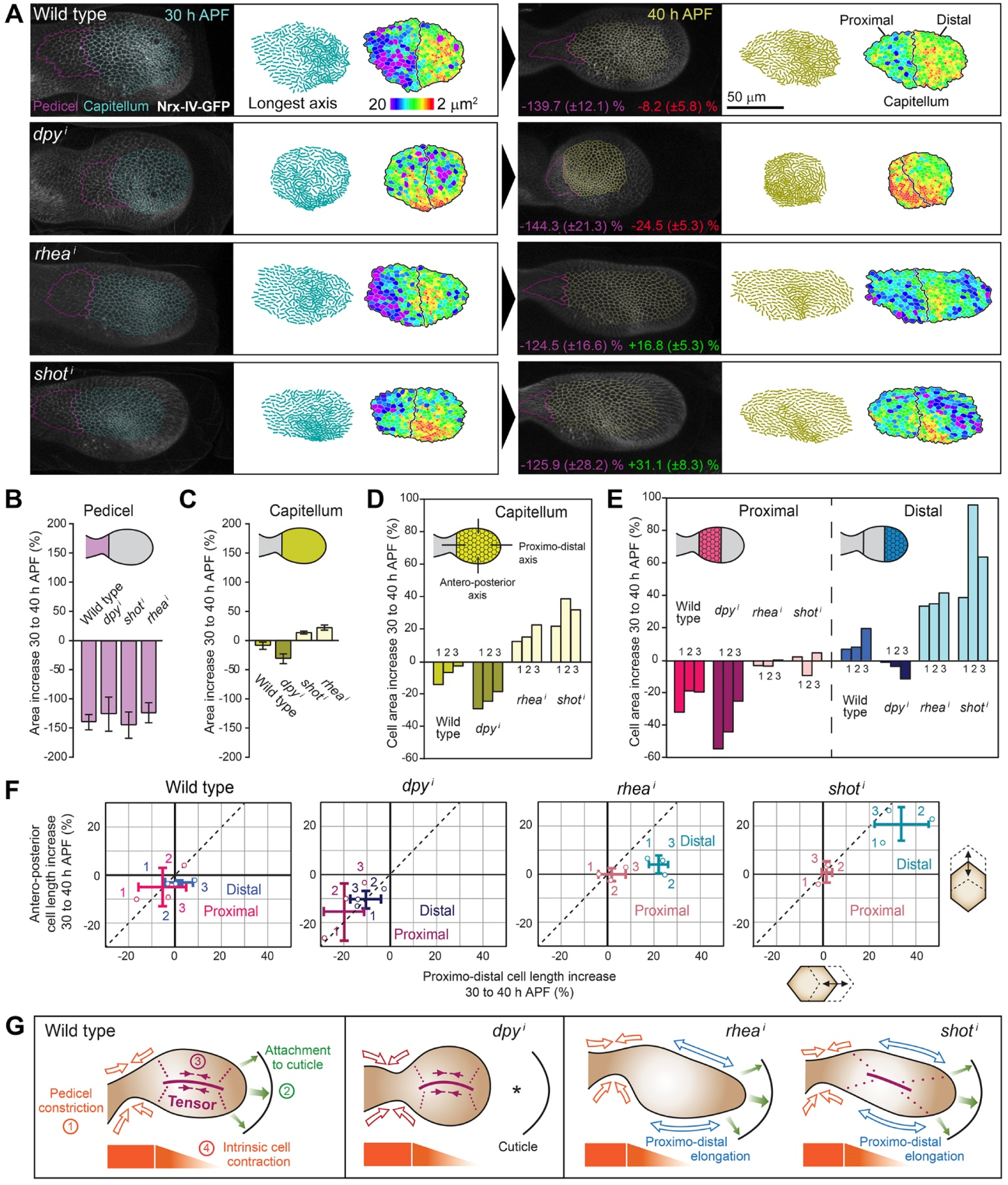
DV coupling counters tissue-elongating tension. (A) Time-lapse imaging (MIP of confocal dorsal xy sections) of haltere development in wild type, *rn>dpy^i^*, *rn>rhea^i^* and *rn>shot^i^* pupae from 30 (left) to 40 h APF (right) (see Suppl. Movie S2). Cell outlines marked with GFP-tagged Neurexin IV (NrxIV^CA06597^-GFP, white). Anterior up, posterior down; proximal left, distal right. A magenta line delimits the same area of the pedicel at 30 and 40 h APF. Transparent cyan and yellow outlines track cells inside the capitellum region. To the right of each image, longest axis of cells (after ellipse fit) and apical cell areas calculated with Icy/Epitools software (Heller et al., 2016) are represented. Two additional movies per genotype were analyzed (see Suppl. Figure S1). Percentages in purple indicate area decrease of the tracked pedicel region ±SD (n=3) from 30 to 40 h APF. Percentages in red (decrease in wild type and *dpy^i^*) or green (increase in *rhea^i^* and *shot^i^*) indicate area change of the tracked capitellum region ±SD (n=3). (B) Total area change of the tracked pedicel regions from 30 to 40 h APF. Error bars represent SD (n=3). (C) Total area change of the tracked capitellum regions. Error bars represent SD (n=3). (D) Changes in average apical cell area inside tracked capitellum region from 30 to 40 h APF. Three movies (1, 2 and 3) were analyzed per genotype. (E) Changes in average apical cell area inside tracked proximal and distal capitellum regions from 30 to 40 h APF. (F) Changes in average antero-posterior and proximo-distal aspects of cells inside tracked proximal and distal capitellum regions from 30 to 40 h APF. Three movies (1, 2 and 3) are represented per genotype. Error bars indicate SD in each axis, crossing at average values (n=3). Dashed lines represent equal change in both axis; note that distal *rhea^i^* and *shot^i^* capitellum regions locate below this line. (G) Forces shaping the haltere, inferred from analysis of time-lapse movies. Pedicel constriction exerts a proximal pull on the organ (1). Due to the attachment to the cuticle, this generates a reaction force in the opposite direction (2), evidenced by the strong retraction of *dpy^i^*halteres. The proximo-distal tension created by these two forces is countered by the DV projections, connecting both surfaces of the haltere to a central tensor (3). When projections are missing (*rhea^i^*) or defective (*shot^i^*), proximo-distal tension is entirely transmitted through the surface, resulting in cell and tissue elongation. In addition to these global forces, a proximo-distal gradient of autonomous cell constriction exists in the capitellum (4).

### Two dorso-ventral bridges remain in the adult haltere

To examine the final stages of haltere development until adulthood, we imaged halteres in vivo at 50, 60, 72, 84 and 96 h APF. We observed that contact between dorsal and ventral surfaces is maintained through late metamorphic stages despite dynamic rearrangements. At 50 h APF a single central DV contact area is still apparent; however, at 60 and 72 h APF this single area has been replaced by four discrete DV contact regions, with three remaining at 84 h APF and two at 96 h APF, near the time of adult eclosion (Figure 7A and B). In the eclosed adult, in fact, these two DV contact regions are preserved: one more central, which we termed the “medial DV bridge”, and the other obliquely positioned in the distal tip of the haltere, which we named the “distal DV bridge” (Figure 7C). Confirming that these structures mediate DV contact in the adult haltere, their dorsal portions, approximately one half, were labeled by dorsal compartment marker *apterous*-GAL4 (*ap*-GAL4) (Figure 7D). In addition, neither the medial nor the distal DV bridges were labeled by posterior compartment marker *hedgehog*-GAL4 (*hh*-GAL4) (Figure 7D), indicating that both structures are formed by cells of the anterior compartment. In all, these data show that two DV bridges remain in the adult haltere, confirming that DV adhesion is not exclusive to the wing; on the contrary, contact between the dorsal and ventral surfaces of the haltere is an essential feature of its developing and final adult forms.

**Figure 7.**
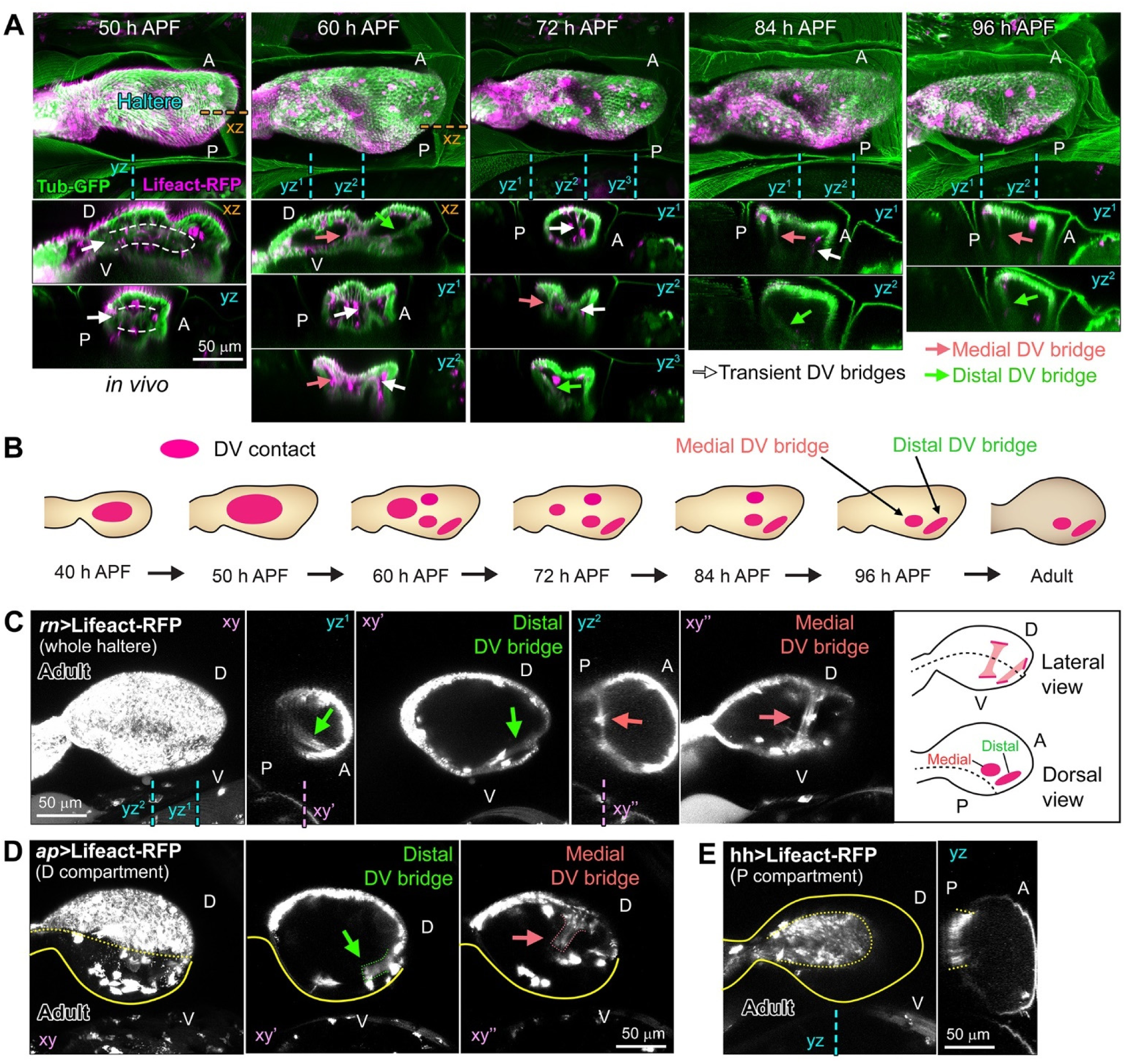
DV contact becomes restricted to two bridges during late haltere development. (A) In vivo images (MIP of confocal dorsal xy sections) of wild type halteres in late metamorphosis (50, 60, 72, 84 and 96 h APF). F-actin and microtubules labeled with *rn-GAL4*-driven expression of Lifeact-RFP (magenta) and Tubulin-GFP (green), respectively. Anterior [A] up, posterior [P] down; proximal left, distal right. Position of xz and yz sections below (dorsal [D] up, ventral [V] down) indicated by discontinuous orange and cyan lines, respectively. Arrows point to DV contacts, including the medial and distal DV bridges remaining in the adult. (B) Schematic drawing of the evolution of DV contact areas during late haltere development. (C) Confocal images of an adult haltere (xy to xy’’: posterior to anterior). Dorsal [D] up, ventral [V] down. Position of yz and xy sections indicated by discontinuous cyan and mauve lines, respectively. F-actin labeled by *rn-GAL4*-driven expression of Lifeact-RFP (white). Arrows point to medial (salmon) and distal (green) DV bridges. Their positions in lateral and dorsal views are depicted in the schematic drawing on the right. (D) Confocal images (xy to xy’’: posterior to anterior) of an adult haltere. Dorsal [D] up, ventral [V] down. Dorsal compartment labeled with *apterous-GAL4-*driven expression of Lifeact-RFP (white). Yellow lines mark DV compartment boundary (dotted) and haltere outline (solid). Arrows point to medial (salmon) and distal (green) bridges; note labeling in their dorsal portions (salmon and green dotted lines). (E) Confocal image of an adult haltere. Dorsal [D] up, ventral [V] down. Posterior compartment labeled with *hedgehog-GAL4*-driven expression of Lifeact-RFP (white). Position of yz section indicated by discontinuous cyan lines. Yellow lines mark AP compartment boundary (dotted) and haltere outline (solid). Note bridges are not labeled.

## DISCUSSION

Here, we investigated how the haltere acquires its characteristic globular shape. Our findings provide critical insights into its metamorphic development and a new context for comparative analysis of wing and haltere morphogenesis. Importantly, our results contradict “hollow haltere” conventional wisdom and the theory that differential BM degradation is responsible for the divergence in shape between the flat wing and the globular haltere (De las Heras et al., 2018; Diaz-de-la-Loza et al., 2018). According to this theory, BM degradation does not take place in the haltere, in contrast with the wing, where BM degradation sets in motion the DV adhesion process. We demonstrate, however, that BM degradation does occur in the haltere as well and is a required step in its morphogenesis. The significance of slightly delayed BM degradation in the haltere (1-2 h after the wing) remains to be determined. Nonetheless, like in the wing, presence of all four major BM components acutely decreases in the haltere during the prepupal period. Furthermore, in both wing and haltere, TIMP overexpression and Mmp2 knockdown resulted in Collagen IV accumulation between the dorsal and ventral surfaces. Also similar to the wing (De las Heras et al., 2018; Diaz-de-la-Loza et al., 2018), preventing BM degradation in the haltere hindered prepupal tissue expansion and gave rise to rounder appendages, further confirming the occurrence of BM degradation in the haltere. Mmp2 expression presents a large peak at the beginning of metamorphosis, highly consistent with hormonal ecdysone control. Other BM degradation processes in metamorphosis such as disc eversion (Srivastava et al., 2007) and fat body dissociation (Jia et al., 2014) are known to require Mmp2 activity as well, suggesting that ecdysone-controlled Mmp2 upregulation is a general mechanism for metamorphic tissue remodeling.

Besides triggering expansion of the wing, prepupal BM degradation precedes wing DV contact and formation of DV projections (Sun et al., 2021). Our results show that resemblance between wing and haltere development extends to these following phases of the morphogenesis of the appendage. Like in the wing, DV projections are found in the haltere. The formation of these projections in the haltere, like in the wing (Sun et al., 2021), requires Laminin, integrin and talin, which concentrate at the area of DV contact, where the basal ends of projections meet. At 40 h APF A single Laminin-rich DV contact area at the center of the haltere capitellum is discernable. Despite later changes to the number and size of DV contact areas, connection between the two haltere surfaces is maintained, including after the adult has eclosed. An additional similarity between haltere and wing uncovered by our study is that Bs (*Drosophila* SRF), a determinant of the DV-adhered wing intervein fate (Fristrom et al., 1994; Montagne et al., 1996), is also expressed in the pupal haltere, and required there for downstream expression of Sdb and normal appendage development. In summary, our results reveal unanticipated similarity in the metamorphic development of halteres and wings, and prove that DV adhesion is not a wing-specific feature but an essential characteristic of haltere development too.

Despite these similarities between the haltere and the wing, our work found key differences in their development, too. Our in vivo imaging showed that both haltere and wing undergo organ compaction between 20 and 24 h APF. Wing projections, however, contract to a larger extent, flattening the appendage by bringing its dorsal and ventral surfaces to a closer distance. It is therefore epithelial height, determined by contraction of DV projections, what distinguishes wing and haltere rather than DV adhesion, found in both. Further research is needed to uncover how this difference in projection contractility is controlled, with many potential Ubx-regulated target genes as candidates to play a role (Choo et al., 2011; Mohit et al., 2006; Pavlopoulos and Akam, 2011; Shlyueva et al., 2016; Slattery et al., 2011; Tomoyasu, 2017). These critical Ubx targets, our findings indicate, are unlikely to be genes coding for MMPs, TIMP or integrins, as previously hypothesized (De las Heras et al., 2018; Diaz-de-la-Loza et al., 2020; Diaz-de-la-Loza et al., 2018), since the difference between haltere and wing is not one of BM degradation or DV adhesion.

A second difference between the haltere and the wing our study underscores is that DV adhesion is critical for their correct development in different periods. In the wing, DV adhesion through tension-resistant projections is required during head eversion to prevent blisters. Our previous research showed that wing blisters originate when fat body adipocytes pushed by increasing hydrostatic pressure penetrate through the hinge at the time of head eversion, and that projections act as a stopper resisting excessive inflation and as a fence preventing fat body entrance (Sun et al., 2021). In the haltere, in contrast, DV adhesion is not required to prevent excessive inflation during head eversion, but later, between 20 and 40 h APF, to preserve organ shape during pedicel contraction. A reason why head eversion is not a critical period for haltere development could be the fact that the haltere does not contract its DV projections after head eversion, and thus trapping of fat body between haltere surfaces may not represent a risk of malformation like in the wing. Nonetheless, we never observed fat body inside mutant halteres, suggesting that the small size of the appendage and narrow pedicel make entrance of adipocytes impossible to begin with.

Once discarded a role for haltere DV adhesion directly equivalent to the one it plays in the wing, our attempt to understand deformations in mutant halteres led us to uncover a complex interplay of forces shaping the organ during metamorphosis. Through in vivo image analysis, we found that the haltere pedicel strongly constricts, pulling the appendage towards proximal. This is reminiscent of the observations of others in the hinge (Aigouy et al., 2010; Etournay et al., 2015), the wing equivalent of the pedicel. Proximal pedicel pull, in turn, is opposed by a reaction force dependent on the attachment of the haltere to the distal cuticle. As a result of these two opposing forces, the tissue is subjected to proximo-distal tension that tends to elongate the haltere. Evidence of the reaction force and resulting tension is the retraction of the organ upon knock down of Dumpy, anchoring the haltere to the cuticle. A requirement for Dumpy in shaping wings and legs during metamorphosis has been reported (Ray et al., 2015). The effect we observe in the haltere is equivalent, as loss of attachment to the cuticle produces proximo-distal shortening of all three appendages. In this context of proximo-distal tension that tends to elongate the haltere, DV connection helps maintain organ shape, as both adult phenotypes and movies reveal. By connecting the surface to a common central structure, DV projections distribute tensile loads and prevent that they are entirely transmitted in-plane. Supporting this, when projections are absent or defective, proximo-distal tension produces organ elongation. In the wing, where hinge contraction is opposed by the attachment of the wing margin to the cuticle (Ray et al., 2015), DV connection could play a role as well in distributing and attenuating tissue-elongating tension. This, however, would be difficult to confirm, since wings lacking DV adhesion or resistant projections are filled with fat body cells after head eversion and deeply malformed by the time of hinge contraction.

In addition to global tissue-wide forces, our study revealed autonomous cell forces at play as well. We found that cell contraction is not restricted to the pedicel region. In the wild type haltere, while distal cells slightly expand their apical area in the 30-40 h APF period, cells in the proximal capitellum region contract. Furthermore, in the absence of cuticle attachment by Dumpy, and thus lack of tissue-elongating tension, cells contract even more in the proximal region and slightly contract in the distal one. These results evidence a pattern of autonomous cell contraction affecting the whole haltere, suggesting a proximo-distal gradient that would include the pedicel as the area of highest contraction. How patterned constriction of the haltere pedicel and wing hinge is achieved is an issue worth further research given the key roles these have in shaping both organs. Moreover, besides tissue-wide strains and graded cell contraction, other cellular processes observed in our movies could significantly contribute to haltere morphogenesis as well, such as cell intercalation, cell proliferation, basal extrusion and cell death, and may also deserve more detailed characterization in the future.

Beyond the analysis of haltere development, finally, our findings add to the repertoire of roles of BM-related matrices in tissue morphogenesis and provide insights into the creation of 3D shape from 2D epithelia. Specifically, we characterized a novel role of an atypical, Laminin-only BM as a critical constituent of a matrix-cytoskeleton tensor required for folding an epithelium into a globular shape. The mechanical properties of the haltere tensor are particularly intriguing, as it contains mainly Laminin, which should lack the tensile strength of Collagen IV (Al-Shaer et al., 2021). The DV connections of haltere and wing, involving partial digestion and fusion of two BMs, are examples of basal-basal linkage matrices (Keeley and Sherwood, 2019). Because of the similar tissue linkage role played by the ECM and its importance in the evolution of propelled motion, DV adheson in the wing and haltere is reminiscent of fin development in fishes, where two epidermal surfaces are joined through their BMs to form the fin apical fold (Carney et al., 2010; Yano et al., 2012; Zhang et al., 2022). Reminiscent also of *Drosophila* appendages, zebrafish mutants for *lama5*, encoding Laminin α5 chain, fail to form an apical fold and display malformed fins (Webb et al., 2007). Regarding the cytoskeletal component of the tensor, our study may offer general lessons on how tissue strains are handled by epithelia to produce curvature. Epithelial force transmission usually occurs in-plane and relies on the zonula adherens connecting neighboring cells. The physical properties of an epithelium, therefore, are largely determined by the elasticity and plastic behavior of these planar intercellular connections and their associated acto-myosin networks. Folding an epithelium into a globular shape, however, will necessarily subject it to pulls and strains in non-planar directions. In this situation, the epithelial tubulin cytoskeleton, containing abundant microtubules oriented apico-basally along lateral membranes, may assume a more critical role in distributing forces to achieve a given organ shape. These microtubules are anchored to the sub-apical and basal actin networks of the zonula adherens and focal adhesion, respectively, ensuring tissue integrity and coordinated strain responses, both passive and active. Metamorphic development of the *Drosophila* haltere, in this light, offers an excellent and simple system for modeling the generation of complex shapes and for dissecting cellular and ECM contributions to 3D morphogenetic force production.

## Acknowledgements

We thank the Bloomington Drosophila Stock Center, Kyoto Drosophila Stock Center, Vienna Drosophila Resource Center, and Tsinghua Fly Center for numerous fly strains, and the Developmental Studies Hybridoma Bank for anti-Mys antibody. This work was funded by grant 32150710524 from the National Natural Science Foundation of China, grant PID2021-122119NB-I00 from Ministerio de Ciencia, Innovación y Universidades and the “Severo Ochoa” Program for Centers of Excellence CEX2021-001165-S. Work in the laboratory of E.S.-H. was funded by grant PID2020-113318GB-I00 from Ministerio de Ciencia, Innovación y Universidades and institutional support from Fundación Ramón Areces to the CBMSO.

## Author contributions

Y.S., P.M. and T.S. conducted experiments. Y. S., E.S.-H. and J.C.P.-P. analyzed the data. J.C.P.-P. wrote the manuscript with input from all authors.

## MATERIALS AND METHODS

### Drosophila genetics

Standard fly husbandry and genetic methods were used to create intermediate strains and calculate segregation of transgenes and mutations in the progeny of crosses. *Drosophila melanogaster* strains and genetic crosses were maintained on standard medium (yeast 24.5 g/L, cornmeal 50 g/L, agar 10 g/L, white granulated sugar 7.25 g/L, brown granulated sugar 30 g/L, propionic acid 4 mL/L, methyl-4-hydroxybenzoate 1.75 g/L and absolute alcohol 17.5 mL/L) and raised at 25°C. The GAL4-UAS binary system was used to drive UAS transgenes and RNAi constructs in halteres and wings under control of *rn-GAL4*. Other drivers were *bs-GAL4, ap-GAL4* and *hh-GAL4*. Penetrance of wing blister phenotypes is reported below representative images in Figure 2I. For fixed tissue samples, images shown are representative of at least 5 animals (10 halteres or 10 wings) of the corresponding genotype and developmental time point. Detailed genotypes in each experiment are listed in Suppl. Table S2. Original strains used were:

*y w; vkgG454.GFP* (DGRC_110692)

*w trol^ZCL1700^.GFP* (DGRC_110807)

*y w; Ndg^fTRG.638^.sGFP* (VDRC_318629)

*y w; LanB1^fTRG.681^.sGFP* (VDRC_318180)

*w; rn-GAL4 / TM3, Sb* (BDSC_7405)

*w; UAS-myr.RFP* (BDSC_7118)

*w; UAS-TIMP.13.1* (gift from Tian Xu, Srivastava et al., 2007)

*w; UAS-Mmp2.RNAi^VDRC.v107888^*(VDRC_107888)

*w; UAS-EcR.RNAi^VDRC.v37058^* (VDRC_37058)

*w; UAS-Mmp2#4* (gift from Tian Xu, Srivastava et al., 2007)

*w^1118^* (BDSC_3605)

*w; UAS-Mmp1.RNAi^VDRC.v330331^*(VDRC_330331)

*w; UAS-Mmp1.RNAi^VDRC.v101505^*(VDRC_101505)

*y sc v; UAS-Mmp1.RNAi^TRIP.JF01336^* (BDSC_31489)

*y sc v; UAS-Mmp2.RNAi^TRIP.HMJ23143^* (BDSC_61309)

*w; UASp-GFPS65C.αTub84B / TM3, Sb* (BDSC_7373)

*w; UAS-Lifeact.RFP* (BDSC_58362)

*y w; rhea^MI00296-mCh.0.mCherry^*(BDSC_39648)

*Ubx^pbx-1^ / T(2;3)apXa* (BDSC_3449)

*Df(3R)Ubx^109^ / Dp(3;3)P5* (BDSC_3486)

*w; UAS-LanB1.RNAi^VDRC.v23121^* (VDRC_23121)

*w; UAS-mys.RNAi^NIG.1560R-1^* (NIG_1560R-1)

*y sc v; UAS-rhea.RNAi^TRiP.HMS00799^* (BDSC_32999)

*y sc v; UAS-shot.RNAi^TRiP.JF02971^* (BDSC_28336)

*y sc v; UAS-Sdb.RNAi^TRiP.HMJ23508^* (BDSC_61925)

*w; bs-GAL4 / CyO* (BDSC_25753)

*w; UAS-bs.RNAi^VDRC.v330226^*(VDRC_330226)

*y sc v; UAS-dpy.RNAi^TRiP.HMS01580^* (BDSC_36691)

*y w; NrxIV^CA06597^.GFP* (BDSC_50798)

*y w; ap-GAL4^MD544^*/ CyO (BDSC_3041)

*w; hh-GAL4 / TM6B* (gift from Isabel Guerrero, Mullor and Guerrero, 2000).

### Imaging of fixed pupal halteres and wings

Pupae were staged at either 0 h APF (white pupae) or 12 h APF (head just everted). For dissection, staged pupae were rinsed in tap water to remove medium and pre-dissected in PBS. Pre-dissection was done by holding the pupa with fine tip forceps and cutting away anterior and posterior ends with scissors. To allow entrance of fixative, we cut pupal epidermis and pupal case along the dorsal midline and removed with forceps and scissors as much fat body, gut and other organs as possible without disrupting epidermis and overall morphology. After pre-dissection, pupae still inside their pupal cases were fixed in 4% PFA for 1 h and washed in PBS (3×10 min). Animals were then taken out of the pupal case (and cuticle, in the case of 40 h APF halteres) and either dissected from the rest of the pupal epidermis on a glass slide inside a drop of DAPI-Vectashield mounting medium (Vector Laboratories, Cat# H-1200) or subjected to antibody or phalloidin staining as explained below. Once mounted, fixed pupal halteres and wings, were imaged in a Zeiss LSM780 upright confocal microscope equipped with Plan-Apochromat 63× oil (NA 1.4), 40× water (NA 1.2) and 20× air (NA 0.8) objectives. The image of a *pbx* homeotically transformed haltere (Figure 4G) was acquired in a Nikon A1R+ inverted confocal microscope using a 40× oil objective (NA 1.3). Maximum intensity projections of xy sections and orthogonal z sections of confocal stacks were obtained with Zeiss Zen software or ImageJ-FIJI.

### Immunohistochemistry

The following antibodies and dyes were used: mouse monoclonal anti-Mys (1:200, Developmental Studies Hybridoma Bank, Cat# CF.6G11), rabbit anti-Sdb (1:50, Sun et al., 2021), donkey anti-mouse IgG conjugated to Alexa-555 (1:200, Thermo Fisher Scientific, Cat# A-31570), donkey anti-rabbit IgG conjugated to Alexa-555 (1:200, Thermo Fisher Scientific, Cat# A-31572) and Phalloidin conjugated to TRITC (1:200, Santa Cruz, Cat# sc-3015309).

For antibody stainings (Figure 2E and Figure 5J and K), 40 h APF pupal samples fixed as described above were removed from the pupal case and cuticle, and washed in PBS (3×10 min). Samples were then blocked in PBT-BSA (PBS containing 0.3% Triton X-100 detergent, 1% BSA, and 250 mM NaCl) and incubated overnight with primary antibody in PBT-BSA at 4°C. Next, samples were washed in PBT-BSA (3×20 min), incubated for 2 h with secondary antibody in PBT-BSA at room temperature, and washed in PBT-BSA (3×20 min) and PBS (3×10 min). Finally, halteres were dissected from the rest of the pupal epidermis on a glass slide inside a drop of DAPI-Vectashield and mounted.

For phalloidin staining (Figure 4G), 40 h APF pupal samples were removed from the pupal case and cuticle, washed in PBT (PBS containing 0.1% Triton X-100 detergent; 3×15 min), and incubated with phalloidin (1:200 in PBT) for two hours at room temperature. Next, samples were washed in PBT (3×15 min) and PBS (3×10 min). Finally, halteres were dissected from the rest of the pupal epidermis on a glass slide inside a drop of DAPI-Vectashield and mounted.

### Imaging of pupal halteres and wings in vivo

Pupae were staged at head eversion stage (12 h APF) and thoroughly washed with tap water and a brush to remove medium. Next, the pupa was dried in filter paper, attached to a glass slide through double-side sticky tape and carefully removed from the pupal case with the help of forceps. The pupa was then placed on a glass bottom dish (Cellvis, Cat# D35-20-1) in which a little drop of halocarbon oil 700 (Sigma, Cat# H8898) had been added, just enough to attach the dorsal side of the wing and a dorso-lateral portion of the pupa through surface tension without spreading to the rest of the animal. In this position, corrected with forceps as needed, the wing is slightly displaced towards anterior, which allows imaging of both the wing and the now exposed haltere while preserving their 3D morphology. Because imaging was performed in an upright microscope, the position of the glass bottom dish must be inverted (pupa is hanging from the glass bottom of the dish, held by surface tension). Using this method, pupae could develop normally until around 70 h APF (orange/red eye color). Imaging was conducted at room temperature (22-23°C) in a Zeiss LSM780 upright confocal microscope using 40× water (NA 1.2) and 20× air (NA 0.8) objectives for the haltere and wing, respectively. For single-time images of halteres (Figure 3A-C, Figure 5A-F and Figure 7A), stacks of 80 to 120 sections spaced 1 μm in the z axis were acquired. For time-lapse imaging of halteres (Figure 4A, Figure 6A and Suppl. Figure S1) and wings (Figure 4B), stacks of 60 to 100 sections spaced 1 μm in the z axis were acquired every 200 seconds. Maximum intensity projections and orthogonal sections of confocal stacks were obtained with Zeiss Zen software.

### Imaging of adult halteres and wings

For confocal imaging of adult halteres (Figure 7C-E), adults were collected approximately 3 hours after eclosion. Their wings and legs were cut off under a stereomicroscope while anesthetized in CO_2_. Next, animals were placed on a glass bottom dish (Cellvis) in which a drop of halocarbon oil 700 (Sigma) had been added, enough to attach the lateral portion of the animal to the glass through surface tension. Because imaging was performed in an upright microscope, the glass bottom dish was inverted (animal is hanging from the glass bottom of the dish). Imaging was conducted in a Zeiss LSM780 upright confocal microscope using a 40× water (NA 1.2) objective.

To image adult halteres in the scanning electron microscope (Figure 2E and Figure 5G-I, M and N), adults were anesthetized with CO_2_ and their legs cut off. Next, the legless animal was gently pressed in a lateral orientation against a piece of electrically conductive adhesive tape, and imaged in a FEI Quanta 200 environmental scanning electron microscope in LowVac mode using large field detector. To image adult wings (Figure 2I), they were cut from animals with scissors and imaged in a Leica M125 stereomicroscope without mounting. The image of a *pbx* homeotically transformed adult haltere (Figure 4F) was taken in a Leica MZ12 stereomicroscope.

### qRT-PCR

To extract RNA from halteres and wings (Figure 2J), approximately 220 halteres and 140 wings per genotype were dissected in PBS. To extract RNA from whole animals (Figure 2K), 10 animals per time point were collected and washed in PBS to remove medium. Samples were then homogenized in TRIzol reagent (Life technologies, Cat# 15596018) with BCP phase separation reagent (Molecular Research Center, Cat# BP 151). Total RNA was extracted and cDNA was synthesized from 2.0 μg (whole animal) or 1 μg (wings and halteres) of RNA using ABScript III RT Master Mix for qPCR with gDNA remover (ABclonal, cat# RK20429). qRT-PCR was performed in a 10 μl reaction system with iTaq™ Universal SYBR® Green Supermix (Bio-Rad, Cat# 172-5121). Each genotype and developmental time point had three technical replicates. The reaction system was composed of 5 μL of 2x Supermix, 0.5 μL of forward primer, 0.5 μL of reverse primer, 1 μL of cDNA, 3 μl of RNase free distilled water. Rp49 expression was used as a reference for normalization. Fold change with respect to the control was calculated with Bio-Rad CFX Manager software. Primers used (Tsingke Biological Technology) were as follows:

MMP2-F: 5’-AACGACGACCGCATGAAGGTG-3’

MMP2-R: 5’- GAAGTGGTTGATCCTTAGCTCCC-3’

RP49-F: 5’- GGCCCAAGATCGTGAAGAAG-3’

RP49-R: 5’- ATTTGTGCGACAGCTTAGCATATC-3’

### Quantification

Signal intensity from fluorescently-tagged BM proteins (Figure 1B-D and Figure 2B, D and G) was measured in ImageJ-FIJI. The BM area in five images per genotype and time point image was delimited with the “Polygon” selection tool and mean intensity inside (intensity divided by area) was calculated with the “Measure” function. Quantification of Sdb expression (Figure 5D) was performed in the same way, delimiting the area of the whole haltere for measurement. Statistical comparison of Sdb expression was performed with GraphPad Prism software. The difference with the wild type was significant (p<0.01=**) in a non-parametric Mann-Whitney test.

For quantification of cell behaviors in pupal halteres and wings from 14 to 26 h APF movies (Suppl. Movie S1), we used the “Line” and “Polygon” selection tools in ImageJ-FIJI to measure (“Measure” function) at 2 h intervals the following parameters in the dorsal haltere central region and six different wing regions, indicated by squares in Figure 4A and B, respectively: apical cell surface area (average values inside each square are plotted in Figure 4C), distance from the apical surface to the basal end (values measured in the center of the corresponding square are plotted in Figure 4D), and distance from the apical dorsal surface to the cuticle (values measured in the center of the corresponding square are plotted in Figure 4D).

For analysis of 30 to 40 h APF haltere movies (Suppl. Movie S1), outlines of cells in the dorsal haltere were manually traced. The boundaries of traced capitellum and pedicel regions and approximate proximo-distal limits were manually tracked through the recordings, whereas outlines of cells inside were drawn only in the initial (30 h APF) and final (40 h APF) frames (Figure 6A and Suppl. Figure S1A-D). To calculate area change in pedicel (Figure 6B) and capitellum (Figure 6C), we used the “Polygon” selection tool in ImageJ-FIJI to measure (“Measure” function) initial and final areas. Calculation and graphic representation of apical areas of individual cells in the capitellum region (Figure 6A, D and E and Suppl. Figure S1A-D) and long axis of their ellipse fits (Figure 6A and Suppl. Figure S1A-D) were performed with Cell Graph Epitools plugins for Icy after skeletonizing initial and final images (Heller et al., 2016). Calculation of antero-posterior and proximo-distal lengths of cells (Figure 6F) was performed on their ellipse fits by using the following formulas:

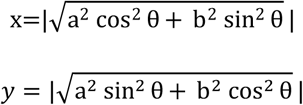

where *x* and *y* are, respectively, the proximo-distal and antero-posterior aspect lengths of the ellipse with major axis length a, minor axis length b and major axis angle θ with respect to the proximo-distal axis. Aspect lengths in other directions (Suppl. Figure S1F) were calculated in an analogous way. Distance from distal tip to the cuticle (Suppl. Figure S1E) was measured with the “Line” selection tool in ImageJ-FIJI.

## Supplemental Materials

**Suppl. Figure S1.**
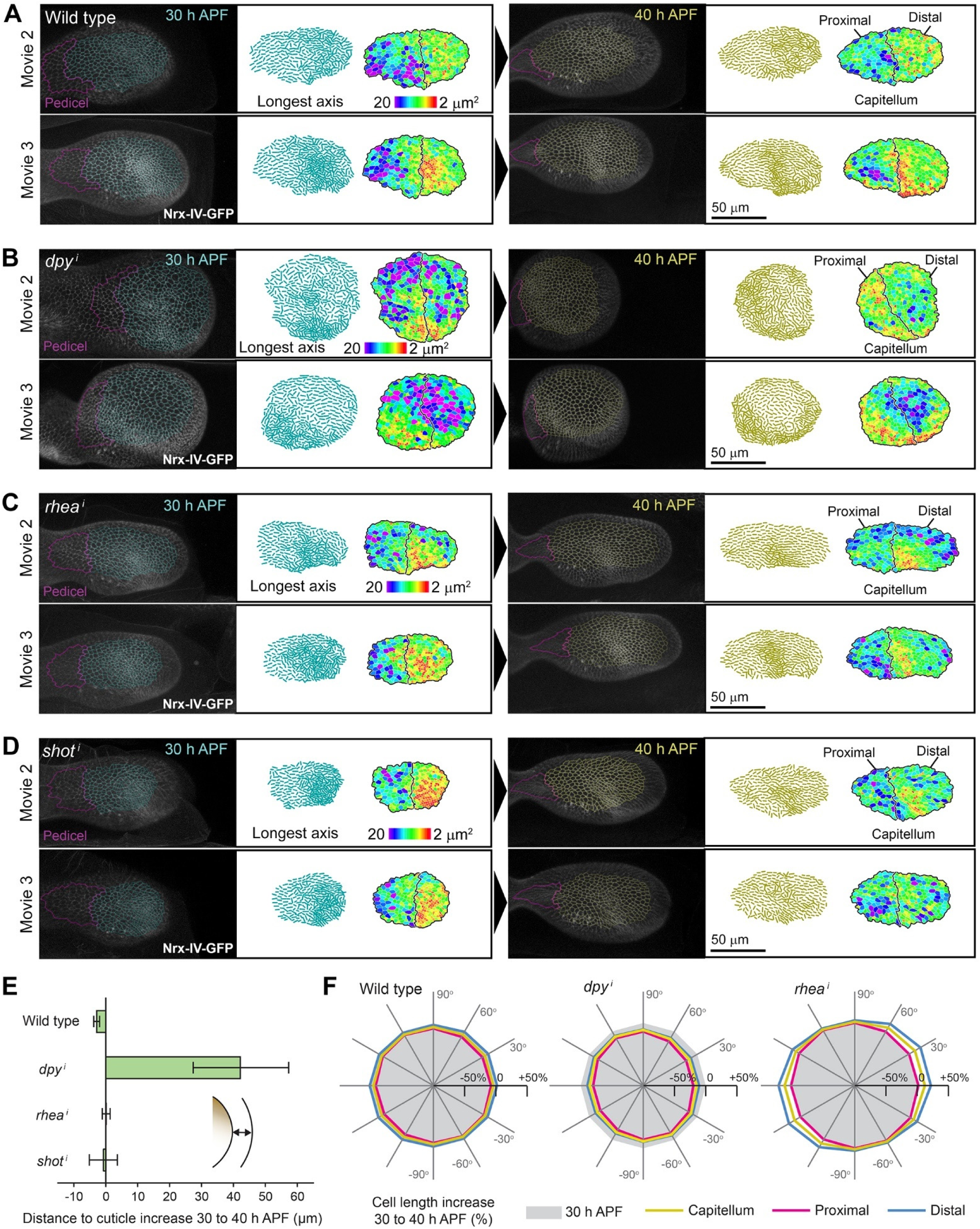
Analysis of 30-40 h APF haltere movies. (A) Time-lapse imaging of pupal haltere development from 30 h APF (left) to 40 h APF (right) in two additional wild type halteres. Cell outlines marked with GFP-tagged, endogenously expressed septate junction protein Neurexin IV (NrxIV^CA06597^-GFP, white). Images are maximum intensity projections (MIP) of confocal xy sections of the haltere dorsal surface. Anterior up, posterior down; proximal left, distal right. A magenta line delimits the same area of the pedicel at 30 h and 40 h APF. Transparent overimposed cell outlines in cyan (30 h APF) and yellow (40 h APF) track the cells inside the same area of the capitellum region. To the right of each image, longest axis of cells (after ellipse fit) and apical cell areas calculated with Icy/Epitools software (B) Time-lapse imaging of pupal haltere development from 30 h APF (left) to 40 h APF (right) in two additional *rn>dpy^i^* halteres. (C) Time-lapse imaging of pupal haltere development from 30 h APF (left) to 40 h APF (right) in two additional *rn>rhea^i^* halteres. (D) Time-lapse imaging of pupal haltere development from 30 h APF (left) to 40 h APF (right) in two additional *rn>shot^i^* halteres. (E) Changes in distance from the distal tip of the haltere to the cuticle from 30 to 40 h APF. Error bars represent SD (n=3). Note large increase in *rn>dpy^i^*. (F) Changes in average aspect of cells inside tracked proximal and distal capitellum regions from 30 to 40 h APF at the indicated directions. 0° (or 180°) and 90° (or −90°) correspond to the proximo-distal and antero-posterior axis, respectively (see Figure 6F).

**Suppl. Table S1.** Defects in wings and halteres upon knock down of *Mmp1* and *Mmp2*. Penetrance of wing and haltere defects is expressed as number of defective appendages/number of appendages examined. Results are shown for three different *Mmp1* transgenic RNAi lines and two *Mmp2* lines.

| Genotype | RNAi line | Wing blistered | Haltere deformed |
| --- | --- | --- | --- |
| *rn-GAL4 > UAS-Mmp1<sup>i</sup>*<br>(*Matrix metalloproteinase 1*) | VDRC SHRNA-330331 | 0 / 22 | 0 / 22 |
|  | VDRC KK-101505 | 0 / 17 | 0 / 17 |
|  | TRIP JF01336 | 0 / 17 | 0 / 17 |
| *rn-GAL4 > UAS-Mmp2<sup>i</sup>*<br>(*Matrix metalloproteinase 2*) | VDRC KK-107888 | 19 / 23 | 16 / 23 (round) |
|  | TRIP HMJ23143 | 6 / 13 | 6 / 13 (round) |

**Suppl. Table S2. Experimental genotypes** (<u>Excel file</u>). Detailed genotypes of animals in all experiments, organized by Figure number.

**Suppl. Movie S1. Behavior of DV projections in haltere and wing: 14 to 26 h APF** (<u>MP4 file</u>). Time-lapse imaging of haltere and wing development from 14 to 26 h APF (=2 to 14 h AHE). Expression of Lifeact-RFP (F-actin, magenta) and Tub-GFP (microtubules, green) driven by *rn-GAL4*. Position of xz and yz sections indicated by discontinuous orange and cyan lines, respectively. Dashed green lines mark the chitinaceous cuticle, visible due to autofluorescence. Arrow points to haltere DV contact.

**Suppl. Movie S2. Haltere morphogenesis: 30 to 40 h APF** (<u>MP4 file</u>). Time-lapse imaging of haltere development in wild type, *rn>dpy^i^*, *rn>rhea^i^* and *rn>shot^i^* pupae from 30 to 40 h APF. Cell outlines marked with GFP-tagged Neurexin IV (NrxIV^CA06597^-GFP, white). Transparent outlines in cyan (30 h APF) and yellow (40 h APF) track cells inside the capitellum region. Dashed grey lines mark the chitinaceous cuticle, visible due to autofluorescence.

## References

Aigouy, B., Farhadifar, R., Staple, D.B., Sagner, A., Röper, J.C., Jülicher, F., and Eaton, S. (2010). Cell Flow Reorients the Axis of Planar Polarity in the Wing Epithelium of Drosophila. Cell 142, 773–786.

Al-Shaer, A., Lyons, A., Ishikawa, Y., Hudson, B.G., Boudko, S.P., and Forde, N.R. (2021). Sequence-dependent mechanics of collagen reflect its structural and functional organization. Biophys J 120, 4013–4028.

Bard, J. (1990). Morphogenesis: The Cellular and Molecular Processes of Developmental Anatomy (Cambridge: Cambridge University Press).

Bender, W., Akam, M., Karch, F., Beachy, P.A., Peifer, M., Spierer, P., Lewis, E.B., and Hogness, D.S. (1983). Molecular Genetics of the Bithorax Complex in Drosophila melanogaster. Science 221, 23–29.

Carney, T.J., Feitosa, N.M., Sonntag, C., Slanchev, K., Kluger, J., Kiyozumi, D., Gebauer, J.M., Coffin Talbot, J., Kimmel, C.B., Sekiguchi, K., et al. (2010). Genetic Analysis of Fin Development in Zebrafish Identifies Furin and Hemicentin1 as Potential Novel Fraser Syndrome Disease Genes. PLoS Genet 6, e1000907.

Choo, S.W., White, R., and Russell, S. (2011). Genome-wide analysis of the binding of the Hox protein Ultrabithorax and the Hox cofactor Homothorax in Drosophila. PLoS One 6, e14778.

Crickmore, M.A., and Mann, R.S. (2006). Hox control of organ size by regulation of morphogen production and mobility. Science 313, 63–68.

Dai, J.L., Ma, M.Q., Feng, Z., and Pastor-Pareja, J.C. (2017). Inter-adipocyte Adhesion and Signaling by Collagen IV Intercellular Concentrations in Drosophila. Curr Biol 27, 2729–2740.e2724.

De las Heras, J.M., Garcia-Cortes, C., Foronda, D., Carlos Pastor-Pareja, J., Shashidhara, L.S., and Sanchez-Herrero, E. (2018). The Drosophila Hox gene Ultrabithorax controls appendage shape by regulating extracellular matrix dynamics. Development 145, dev161844.

de Navas, L.F., Garaulet, D.L., and Sanchez-Herrero, E. (2006). The Ultrabithorax Hox gene of Drosophila controls haltere size by regulating the Dpp pathway. Development 133, 4495–4506.

Diaz-de-la-Loza, M.-d.-C., Loker, R., Mann, R.S., and Thompson, B.J. (2020). Control of tissue morphogenesis by the HOX gene Ultrabithorax. Development 147, dev184564.

Diaz-de-la-Loza, M.-d.-C., Ray, R.P., Ganguly, P.S., Alt, S., Davis, J.R., Hoppe, A., Tapon, N., Salbreux, G., and Thompson, B.J. (2018). Apical and Basal Matrix Remodeling Control Epithelial Morphogenesis. Dev Cell 46, 23–39.e25.

Dickerson, B.H., de Souza, A.M., Huda, A., and Dickinson, M.H. (2019). Flies Regulate Wing Motion via Active Control of a Dual-Function Gyroscope. Curr Biol 29, 3517–3524.e3513.

Etournay, R., Popovic, M., Merkel, M., Nandi, A., Blasse, C., Aigouy, B., Brandi, H., Myers, G., Salbreux, G., Jülicher, F., et al. (2015). Interplay of cell dynamics and epithelial tension during morphogenesis of the Drosophila pupal wing. Elife 4, e07090.

Fristrom, D. (1988). The cellular basis of epithelial morphogenesis. A review. Tissue Cell 20, 645–690.

Fristrom, D., Gotwals, P., Eaton, S., Kornberg, T.B., Sturtevant, M., Bier, E., and Fristrom, J.W. (1994). blistered: a gene required for vein/intervein formation in wings of Drosophila. Development 120, 2661–2671.

Fristrom, D., Wilcox, M., and Fristrom, J. (1993). The distribution of PS integrins, laminin A and F-actin during key stages in Drosophila wing development. Development 117, 509–523.

Godenschwege, T.A., Pohar, N., Buchner, S., and Buchner, E. (2000). Inflated wings, tissue autolysis and early death in tissue inhibitor of metalloproteinases mutants of Drosophila. Eur J Cell Biol 79, 495–501.

Goodwin, K., Ellis, S.J., Lostchuck, E., Zulueta-Coarasa, T., Fernandez-Gonzalez, R., and Tanentzapf, G. (2016). Basal Cell-Extracellular Matrix Adhesion Regulates Force Transmission during Tissue Morphogenesis. Dev Cell 39, 611–625.

Guillot, C., and Lecuit, T. (2013). Mechanics of Epithelial Tissue Homeostasis and Morphogenesis. Science 340, 1185–1189.

Haigo, S.L., and Bilder, D. (2011). Global Tissue Revolutions in a Morphogenetic Movement Controlling Elongation. Science 331, 1071–1074.

Harmansa, S., Erlich, A., Eloy, C., Zurlo, G., and Lecuit, T. (2023). Growth anisotropy of the extracellular matrix shapes a developing organ. Nat Commun 14, 1220.

Heisenberg, C.P., and Bellaiche, Y. (2013). Forces in Tissue Morphogenesis and Patterning. Cell 153, 948–962.

Heller, D., Hoppe, A., Restrepo, S., Gatti, L., Tournier, A.L., Tapon, N., Basler, K., and Mao, Y. (2016). EpiTools: An Open-Source Image Analysis Toolkit for Quantifying Epithelial Growth Dynamics. Dev Cell 36, 103–116.

Henchcliffe, C., Garcia-Alonso, L., Tang, J., and Goodman, C.S. (1993). Genetic analysis of laminin A reveals diverse functions during morphogenesis in Drosophila. Development 118, 325–337.

Hiraiwa, S., Takeshita, S., Terano, T., Hayashi, R., Suzuki, K., Tajiri, R., and Kojima, T. (2024). Unveiling the cell dynamics during the final shape formation of the tarsus in Drosophila adult leg by live imaging. Dev Genes Evol.

Isabella, A.J., and Horne-Badovinac, S. (2015). Building from the Ground up: Basement Membranes in Drosophila Development. In Basement Membranes, J.H. Miner, ed., pp. 305–336.

Jayadev, R., and Sherwood, D.R. (2017). Basement membranes. Curr Biol 27, R207–R211.

Jia, Q.Q., Liu, Y., Liu, H.H., and Li, S. (2014). Mmp1 and Mmp2 cooperatively induce Drosophila fat body cell dissociation with distinct roles. Sci Rep 4, 7535.

Keeley, D.P., and Sherwood, D.R. (2019). Tissue linkage through adjoining basement membranes: The long and the short term of it. Matrix Biol 75-76, 58–71.

Khadilkar, R.J., Ho, K.Y.L., Venkatesh, B., and Tanentzapf, G. (2020). Integrins Modulate Extracellular Matrix Organization to Control Cell Signaling during Hematopoiesis. Curr Biol 30, 3316–3329.e3315.

Khalilgharibi, N., and Mao, Y. (2021). To form and function: on the role of basement membrane mechanics in tissue development, homeostasis and disease. Open Biol 11, 200360.

Klussmann-Fricke, B.-J., Martin-Bermudo, M.D., and Llimargas, M. (2022). The basement membrane controls size and integrity of the Drosophila tracheal tubes. Cell Rep 39.

Lecuit, T., Lenne, P.F., and Munro, E. (2011). Force generation, transmission, and integration during cell and tissue morphogenesis. Annu Rev Cell Dev Biol 27, 157–184.

Lewis, E.B. (1978). A gene complex controlling segmentation in Drosophila. Nature 276, 565–570.

Leys, S.P., and Riesgo, A. (2012). Epithelia, an Evolutionary Novelty of Metazoans. J Exp Zool B Mol Dev Evol 318, 438–447.

Ma, M.Q., Cao, X.Y., Dai, J.L., and Pastor-Pareja, J.C. (2017). Basement Membrane Manipulation in Drosophila Wing Discs Affects Dpp Retention but Not Growth Mechanoregulation. Dev Cell 42, 97–106.e104.

Martin, D., Zusman, S., Li, X.T., Williams, E.L., Khare, N., DaRocha, S., Chiquet-Ehrismann, R., and Baumgartner, S. (1999). wing blister, a new Drosophila laminin alpha chain required for cell adhesion and migration during embryonic and imaginal development. J Cell Biol 145, 191–201.

Matamoro-Vidal, A., Cumming, T., Davidović, A., Levillayer, F., and Levayer, R. (2024). Patterned apoptosis has an instructive role for local growth and tissue shape regulation in a fast-growing epithelium. Curr Biol 34, 376–388.e377.

Matsubayashi, Y., Sánchez-Sánchez, B.J., Marcotti, S., Serna-Morales, E., Dragu, A., Díaz-de-la-Loza, M.D., Vizcay-Barrena, G., Fleck, R.A., and Stramer, B.M. (2020). Rapid Homeostatic Turnover of Embryonic ECM during Tissue Morphogenesis. Dev Cell 54, 33–42.e39.

Mogensen, M.M., and Tucker, J.B. (1988). Intermicrotubular actin-filaments in the transalar cytoskeletal arrays of Drosophila. J Cell Sci 91, 431–438.

Mohit, P., Makhijani, K., Madhavi, M.B., Bharathi, V., Lal, A., Sirdesai, G., Reddy, V.R., Ramesh, P., Kannan, R., Dhawan, J., et al. (2006). Modulation of AP and DV signaling pathways by the homeotic gene Ultrabithorax during haltere development in Drosophila. Dev Biol 291, 356–367.

Montagne, J., Groppe, J., Guillemin, K., Krasnow, M.A., Gehring, W.J., and Affolter, M. (1996). The Drosophila Serum Response Factor gene is required for the formation of intervein tissue of the wing and is allelic to blistered. Development 122, 2589–2597.

Mullor, J.L., and Guerrero, I. (2000). A gain-of-function mutant of patched dissects different responses to the Hedgehog gradient. Dev Biol 228, 211–224.

Page-McCaw, A., Ewald, A.J., and Werb, Z. (2007). Matrix metalloproteinases and the regulation of tissue remodelling. Nat Rev Mol Cell Biol 8, 221–233.

Page-McCaw, A., Serano, J., Sante, J.M., and Rubin, G.M. (2003). Drosophila matrix metalloproteinases are required for tissue remodeling, but not embryonic development. Dev Cell 4, 95–106.

Pastor-Pareja, J.C. (2020). Atypical basement membranes and basement membrane diversity - what is normal anyway? J Cell Sci 133, jcs241794.

Pastor-Pareja, J.C., and Xu, T. (2011). Shaping Cells and Organs in Drosophila by Opposing Roles of Fat Body-Secreted Collagen IV and Perlecan. Dev Cell 21, 245–256.

Pavlopoulos, A., and Akam, M. (2011). Hox gene Ultrabithorax regulates distinct sets of target genes at successive stages of Drosophila haltere morphogenesis. Proc Natl Acad Sci U S A 108, 2855–2860.

Prout, M., Damania, Z., Soong, J., Fristrom, D., and Fristrom, J.W. (1997). Autosomal mutations affecting adhesion between wing surfaces in Drosophila melanogaster. Genetics 146, 275–285.

Ramos-Lewis, W., and Page-McCaw, A. (2019). Basement membrane mechanics shape development: Lessons from the fly. Matrix Biol 75-76, 72–81.

Ray, R.P., Matamoro-Vidal, A., Ribeiro, P.S., Tapon, N., Houle, D., Salazar-Ciudad, I., and Thompson, B.J. (2015). Patterned Anchorage to the Apical Extracellular Matrix Defines Tissue Shape in the Developing Appendages of Drosophila. Dev Cell 34, 310–322.

Ready, D.F., and Chang, H.C. (2023). Interommatidial cells build a tensile collagen network during Drosophila retinal morphogenesis. Curr Biol 33, 2223–2234.e2223.

Roch, F., and Akam, M. (2000). Ultrabithorax and the control of cell morphology in Drosophila halteres. Development 127, 97–107.

Schöck, F., and Perrimon, N. (2002). Molecular Mechanisms of Epithelial Morphogenesis. Ann Rev Cell Dev Biol 18, 463–493.

Sekiguchi, R., and Yamada, K.M. (2018). Basement Membranes in Development and Disease. Curr Top Dev Biol 130, 143–191.

Shlyueva, D., Meireles-Filho, A.C., Pagani, M., and Stark, A. (2016). Genome-Wide Ultrabithorax Binding Analysis Reveals Highly Targeted Genomic Loci at Developmental Regulators and a Potential Connection to Polycomb-Mediated Regulation. PLoS One 11, e0161997.

Singh, A., Thale, S., Leibner, T., Lamparter, L., Ricker, A., Nüsse, H., Klingauf, J., Galic, M., Ohlberger, M., and Matis, M. (2024). Dynamic interplay of microtubule and actomyosin forces drive tissue extension. Nat Commun 15, 3198.

Slattery, M., Ma, L., Négre, N., White, K.P., and Mann, R.S. (2011). Genome-Wide Tissue-Specific Occupancy of the Hox Protein Ultrabithorax and Hox Cofactor Homothorax in Drosophila. PLoS ONE 6, e14686.

Srivastava, A., Pastor-Pareja, J.C., Igaki, T., Pagliarini, R., and Xu, T. (2007). Basement membrane remodeling is essential for Drosophila disc eversion and tumor invasion. Proc Natl Acad Sci U S A 104, 2721–2726.

Sun, T.H., Song, Y.L., Dai, J.L., Mao, D.C., Ma, M.Q., Ni, J.Q., Liang, X., and Pastor-Pareja, J.C. (2019). Spectraplakin Shot Maintains Perinuclear Microtubule Organization in Drosophila Polyploid Cells. Dev Cell 49, 731-+.

Sun, T.H., Song, Y.Z., Teng, D.Q., Chen, Y.A., Dai, J.L., Ma, M.Q., Zhang, W., and Pastor-Pareja, J.C. (2021). Atypical laminin spots and pull-generated microtubule-actin projections mediate Drosophila wing adhesion. Cell Rep 36.

Tomoyasu, Y. (2017). Ultrabithorax and the evolution of insect forewing/hindwing differentiation. Curr Opin Insect Sci 19, 8–15.

Tran, N.V., Montanari, M.P., Gui, J., Lubenets, D., Fischbach, L.L., Antson, H., Huang, Y., Brutus, E., Okada, Y., Ishimoto, Y., et al. (2024). Programmed disassembly of a microtubule-based membrane protrusion network coordinates 3D epithelial morphogenesis in Drosophila. EMBO J 43, 568–594.

Tsuboi, A., Fujimoto, K., and Kondo, T. (2023). Spatiotemporal remodeling of extracellular matrix orients epithelial sheet folding. Sci Adv 9, eadh2154.

Waddington, C.H. (1940). The genetic control of wing development inDrosophila. J Genet 41, 75–113.

Walsh, E.P., and Brown, N.H. (1998). A screen to identify Drosophila genes required for integrin-mediated adhesion. Genetics 150, 791–805.

Walther, R.F., Lancaster, C., Burden, J.J., and Pichaud, F. (2024). A dystroglycan–laminin–integrin axis coordinates cell shape remodeling in the developing Drosophila retina. PLoS Biology 22, e3002783.

Weatherbee, S.D., Halder, G., Kim, J., Hudson, A., and Carroll, S. (1998). Ultrabithorax regulates genes at several levels of the wing-patterning hierarchy to shape the development of the Drosophila haltere. Genes Dev 12, 1474–1482.

Webb, A.E., Sanderford, J., Frank, D., Talbot, W.S., Driever, W., and Kimelman, D. (2007). Laminin α5 is essential for the formation of the zebrafish fins. Dev Biol 311, 369–382.

Wilkin, M.B., Becker, M.N., Mulvey, D., Phan, I., Chao, A., Cooper, K., Chung, H.J., Campbell, I.D., Baron, M., and MacIntyre, R. (2000). Drosophila Dumpy is a gigantic extracellular protein required to maintain tension at epidermal-cuticle attachment sites. Curr Biol 10, 559–567.

Wu, D., Yamada, K.M., and Wang, S. (2023). Tissue Morphogenesis Through Dynamic Cell and Matrix Interactions. Ann Rev Cell Dev Biol 39, 123–144.

Yamanaka, N., Rewitz, K.F., and O’Connor, M.B. (2013). Ecdysone control of developmental transitions: lessons from Drosophila research. Ann Rev Entomol 58, 497–516.

Yano, T., Abe, G., Yokoyama, H., Kawakami, K., and Tamura, K. (2012). Mechanism of pectoral fin outgrowth in zebrafish development. Development 139, 2916–2925.

Yurchenco, P.D. (2011). Basement membranes: cell scaffoldings and signaling platforms. Cold Spring Harb Perspect Biol 3.

Zhang, J.L., Richetti, S., Ramezani, T., Welcker, D., Lütke, S., Pogoda, H.M., Hatzold, J., Zaucke, F., Keene, D.R., Bloch, W., et al. (2022). Vertebrate extracellular matrix protein hemicentin-1 interacts physically and genetically with basement membrane protein nidogen-2. Matrix Biol 112, 132–154.

